# Co-transcriptional Phase Separation of Nucleic Acids at Membrane Surfaces

**DOI:** 10.64898/2026.05.01.721969

**Authors:** Adam Mamot, Thuy An Nguyen, Yusuf Qutbuddin, Svetozar Gavrilovic, Viktoriia Belousova, Souvik Basak, Jan-Hagen Krohn, Nastasja Kaletta, Petra Schwille

**Affiliations:** Department of Cellular and Molecular Biophysics, Max Planck Institute of Biochemistry, Martinsried, Germany; Imaging Center Essen Light Microscopy Unit, University Hospital Essen, Essen, Germany

**Author notes:** Correspondence and requests for materials should be addressed to Adam Mamot and Petra Schwille;.

## Abstract

Transcription is usually framed as information transfer, yet it also injects a new polymer into a crowded, confined environment. Here we demonstrate how spatial confinement to surfaces in a minimal membrane-bound transcription (MBT) system displays the physical consequences of RNA synthesis. Within a dense membrane-tethered DNA network, transcription drives co-transcriptional RNA phase separation: nascent RNA oligomerizes, gels and demixes from a surrounding fluid DNA phase, generating stable spatial patterns while mechanically remodeling the DNA layer. RNA gelation sequesters T7 RNA polymerase, whereas RNA-binding and translation-associated factors reverse gelation and restore fluidity. Thus, in the absence of downstream regulatory machinery, transcription under confinement is sufficient to trigger RNA condensation and nucleic-acid phase separation. The membrane as confining interface catalyzes the onset of DNA-RNA demixing and modulates the morphology of the resulting patterns. Since such large-scale spatial unmixing may be detrimental to cellular physiology, we suggest that one fundamental role of translation is to actively prevent condensation effects created by continuous RNA production.

## Introduction

Transcription is commonly described as a process of information transfer, in which genetic sequences are read, copied, and translated into functional outputs. Yet this description obscures a central physical reality: transcription is not only an informational operation, but also a material one. Each transcription event generates long, highly charged polymer chains within a confined and crowded environment, continuously adding material and perturbing local organization. (1-3) Over evolutionary timescales, cells have therefore faced not only the challenge of managing information flow, but also of accommodating the persistent material flux generated by RNA production. How this flux is buffered, redirected, or suppressed remains a fundamental and incompletely understood problem.

Both DNA and RNA are now widely recognized as active physical components of the cell rather than passive carriers of genetic code. (3, 4) Genomic DNA undergoes continual compaction, looping, and large-scale reorganization driven by its polymeric properties and by active biochemical processes. (4, 5) RNA, in turn, can condense or aggregate with functional consequences in gene regulation, stress responses, and disease. (6, 7) These behaviors do not occur in isolation. Lipid membranes and associated interfaces impose confinement, electrostatic interactions, and mechanical constraints that can strongly influence nucleic-acid organization. (8-14) In modern cells, the nuclear envelope and other spatially organizing nuclear structures define a restricted environment in which RNA synthesis unfolds, linking transcription to a crowded and mechanically constrained system. (1, 15)

Consistent with this view, accumulating evidence indicates that transcription actively participates in shaping genome organization. Transcriptional activity correlates with chromatin decompaction, domain formation, and changes in long-range chromosomal interactions, while RNA itself can contribute to the assembly and maintenance of transcriptionally active compartments. (1-3, 16) However, a key mechanistic question remains unresolved: to what extent do these organizational effects arise from evolved regulatory networks, and to what extent do they reflect unavoidable physical consequences of RNA production itself. In particular, whether sustained RNA synthesis alone—independent of downstream processing, degradation, and regulatory feedback—can drive large-scale reorganization or instability of nucleic-acid structures remains unclear.

Addressing this question directly in living cells is challenging. Cellular transcription is embedded within dense layers of regulatory proteins, chromatin modifiers, RNA processing pathways, and degradation machinery that continuously buffer physical perturbations and maintain homeostasis. (5) While these mechanisms are essential for cellular viability, they also obscure the primary material effects of transcriptional flux, making it difficult to disentangle cause from compensation. From an evolutionary perspective, this buffering raises a deeper question: are complex gene-regulatory architectures solely required for informational control, or did they evolve in part to suppress intrinsic physical instabilities associated with RNA production?

Minimal experimental systems therefore offer a powerful complementary approach. (17-21) By reducing biological complexity to a defined set of components, such systems allow physical constraints and emergent behaviors to be revealed directly, without presupposing regulatory control. In doing so, they can expose regimes of instability or failure—behaviors that are actively suppressed in vivo, yet nonetheless define fundamental physical limits of gene expression. Such limits are not only relevant for understanding modern cells, but also for reconstructing early gene-expression systems in origins-of-life contexts and for guiding the design of synthetic cells capable of sustained transcription. (8)

To isolate the material consequences of RNA production, an experimental system must satisfy three criteria: it must support sustained transcription, impose defined spatial confinement, and permit direct coupling between RNA synthesis and nucleic-acid organization. Existing approaches typically capture only parts of this design space: cellular systems preserve buffering and regulation, many bulk cell-free systems forgo cell-like confinement, and minimal surface-based systems reconstitute spatially organized gene expression without recreating a lipid-defined boundary.

Here we address this gap using a minimal membrane-bound transcription (MBT) system. In this system, transcription is reconstituted within a lipid-bounded environment that encloses a defined DNA network, creating controlled confinement and well-defined physicochemical boundary conditions. RNA synthesis proceeds continuously in the absence of downstream RNA processing and degradation pathways, allowing the physical consequences of transcriptional flux to be examined directly. By coupling sustained RNA production to a membrane-defined architecture, the MBT system provides a minimal platform to probe how transcription itself can reorganize nucleic acids and generate emergent physical behavior.

## Results

### Transcription within DNA network drives RNA phase separation

We assembled the MBT system by deposition of gene-coding double-stranded DNA molecules onto a supported lipid bilayer (SLB) in the presence of ribonucleotide triphosphates and transcription buffer (Figure 1a). The DNA was site-selectively modified with a 5′-cholesterol moiety on the coding strand and a 5′-Atto647N moiety on the template strand to facilitate membrane binding and detection, respectively. The transcription reaction was initiated by addition of T7 RNA polymerase to a pre-equilibrated system. By spiking the lipid membrane with a fluorescently labeled lipid (Atto488-DOPE) and nascent mRNA with a fluorescent uridine triphosphate analogue (Cy3B-UTP), we monitored membrane-bound transcription by three-color total internal reflection fluorescence (TIRF) microscopy (Figure 1a). In this way, we selectively monitored the system within the DNA layer (evanescent-field penetration depth ∼300 nm for the 642 nm laser).

**Figure 1.**
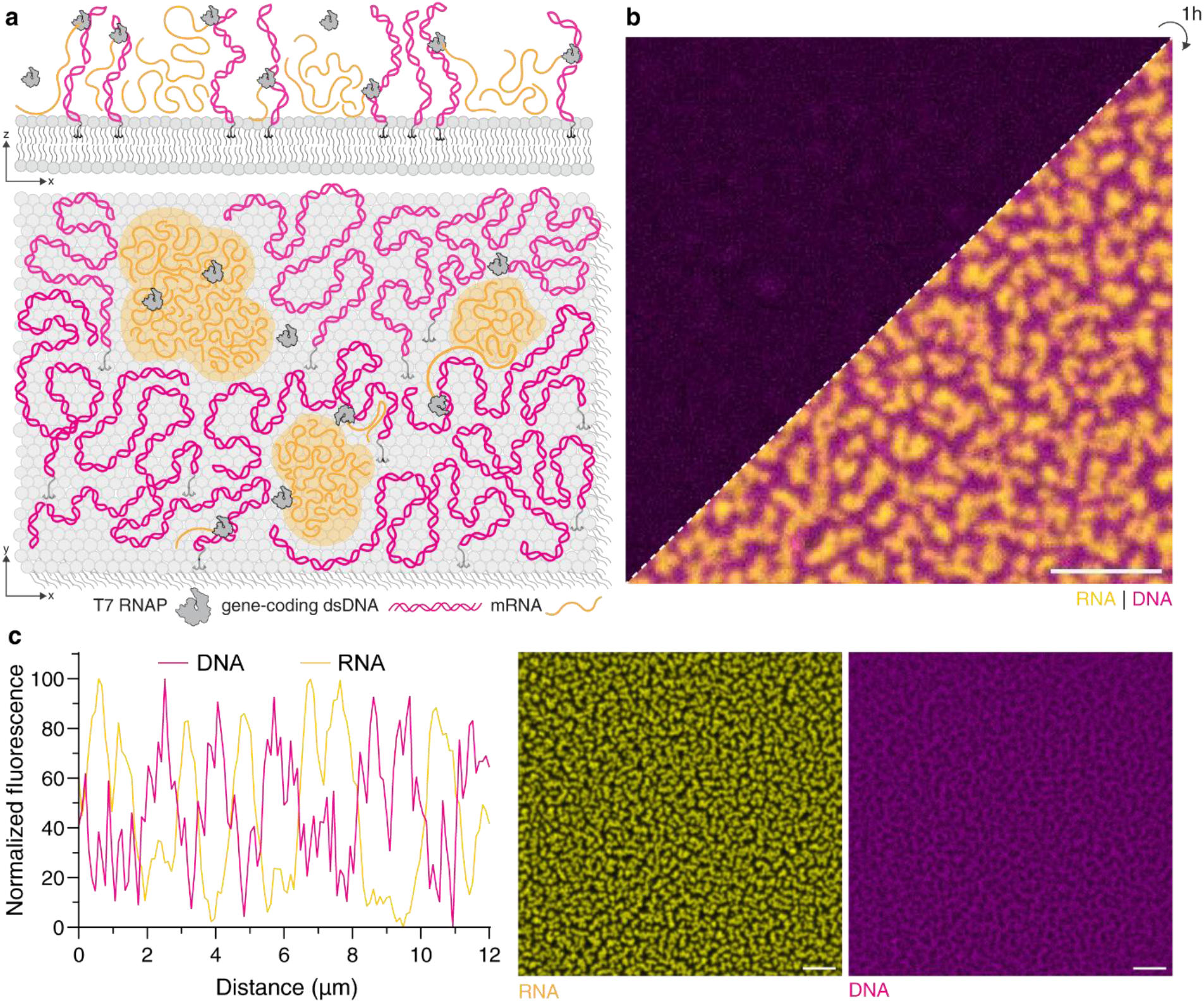
Phase separation of RNA upon transcription in membrane-tethered DNA environment. **a** Schematic representation of the membrane-bound transcription (MBT) set-up on a supported lipid bilayer. Gene-coding DNA was modified with a 5′-cholesterol moiety on the coding strand and a 5′-fluorophore (Atto647N) on the template strand to facilitate membrane binding and detection, respectively. The lipid bilayer was doped with a fluorescently labeled lipid (Atto488-DOPE, cyan) and nascent mRNA with a fluorescent uridine triphosphate analogue (Cy3B-UTP, yellow). The transcription reaction was initiated by addition of T7 RNA polymerase to a pre-equilibrated system. **b** TIRF images of the MBT set-up at the onset of and after 30 min of transcription, showing labelled DNA (magenta), nucleotide/RNA (yellow), and lipid (cyan). **c** DNA and RNA channels (right) and fluorescence intensity profiles over a representative region (left) of a phase-separated system after 30 min of transcription. Scale bars: 10 µm.

To our surprise, we observed the rapid emergence of spatial RNA patterns following transcription initiation. Within approximately 30 minutes, RNA pattern formation reached completion and was accompanied by pronounced anti-localization of DNA and RNA (Figure 1b–c). This behavior was unique to the MBT system: neither transcription in solution nor membrane-tethered DNA in the absence of transcription produced comparable patterns (Figure S1a). Both active transcription and membrane confinement were required, and the phenomenon was independent of RNA labeling and observed across the entire membrane surface (Figure S1b–c).

### Confinement within DNA phase promotes the RNA gelation and liquid-solid phase exclusion

To determine whether RNA patterning at the membrane surface emerges as a consequence of its microenvironment, we first assessed the 3D distribution of RNA. To that end, we recorded DNA and RNA fluorescence intensities as a function of distance from the SLB using confocal microscopy. We found that RNA was enriched approximately twofold within the DNA phase directly at the membrane (Figure 2a). At the same time, overall transcription yield remained unaffected by membrane anchoring, as determined by comparison of isolated RNA yield (Figure 2b). Atomic force microscopy (AFM) revealed that RNA-rich regions were thicker and mechanically stiffer than the surrounding DNA, thus appearing as solid-like islands embedded in a fluid DNA matrix (Figure 2c). Fluorescence recovery after photobleaching (FRAP) confirmed this distinction: RNA-rich regions exhibited negligible recovery, consistent with a gel state, whereas DNA remained fluid both before and after pattern formation (Figure 2d).

**Figure 2.**
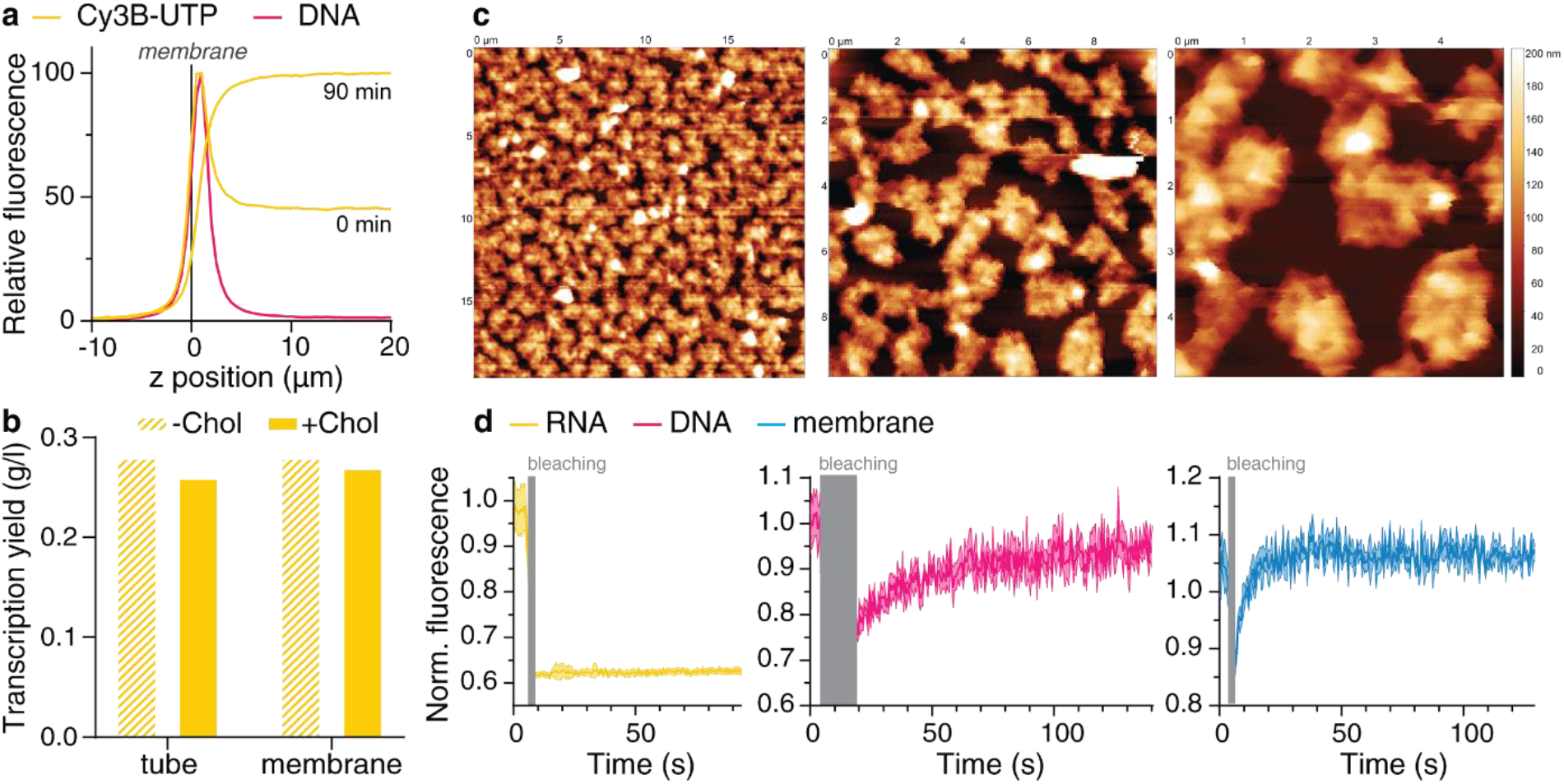
RNA and DNA separate into physically distinct phases in the MBT system. **a** Fluorescence intensity profiles for the membrane, DNA, and RNA channels as a function of the distance (in z) from the lipid membrane, recorded with confocal microscopy. **b** Isolated RNA yields for transcription performed in a test tube and in a membrane chamber, with unmodified or cholesterol-modified, membrane-tethered DNA. The MBT set-up does not influence RNA yield in comparison to conventional *in vitro* transcription in a tube, but results in up-concentration of RNA at the membrane surface. **c** Atomic force microscopy (AFM) images demonstrating the relative height and stiffness of RNA and DNA phases. **d** Fluorescence recovery after photobleaching (FRAP) curves recorded for RNA, DNA, and the membrane after phase separation. Nascent RNA condenses into gel-like structures within the fluid DNA layer.

Time-resolved imaging showed that RNA initially emerged as discrete particles within the DNA network, which subsequently merged into larger structures (Figure 3a). Number-and-brightness (N&B) analysis revealed increasing molecular brightness over time, indicating progressive oligomerization of RNA molecules (Figure 3b). Corresponding number-density maps showed growth from nucleation sites, consistent with self-aggregation and outward expansion of RNA condensates.

**Figure 3.**
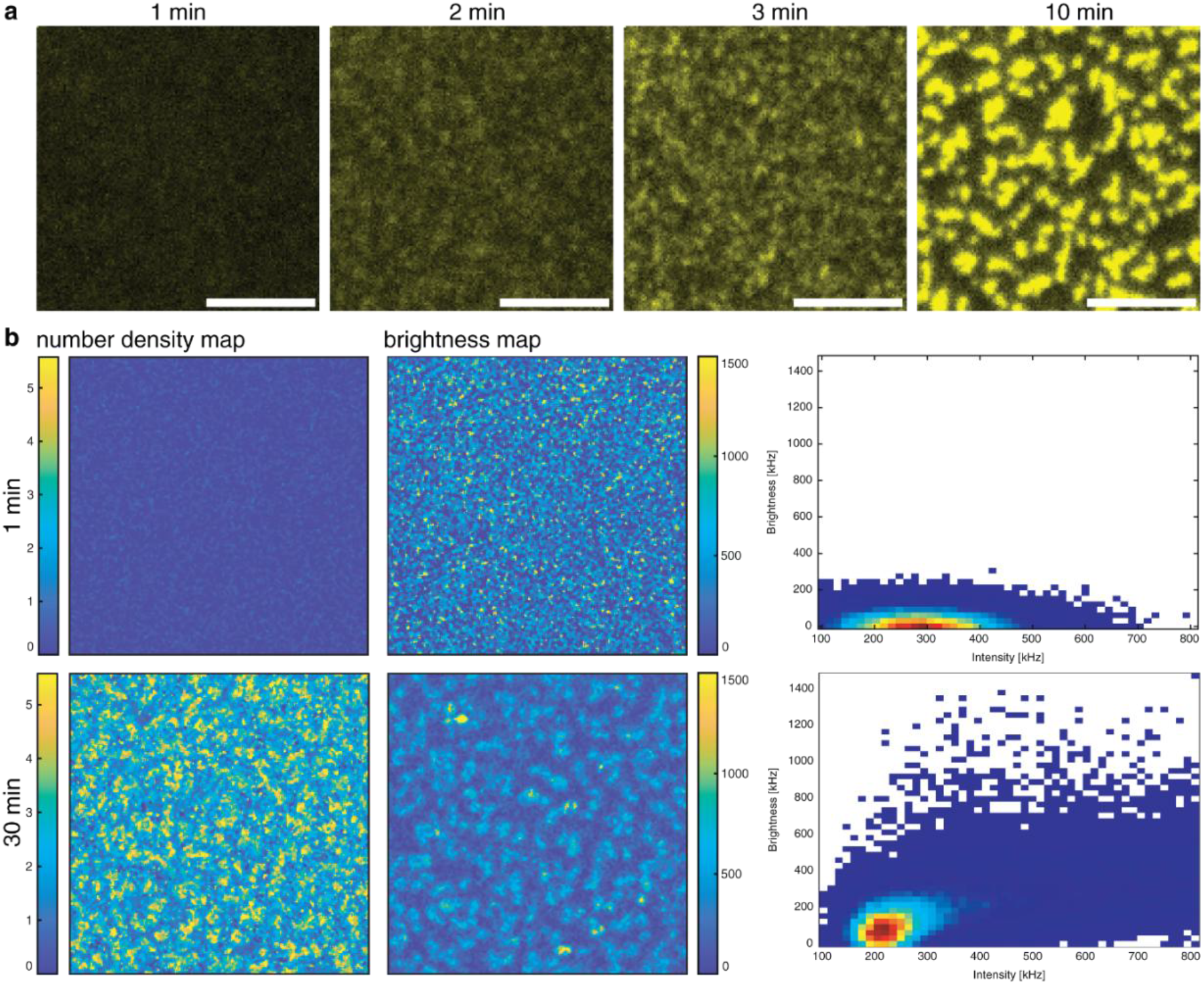
RNA aggregates *via* a liquid-to-solid transition. **a** TIRF images recorded at 1, 2, 3, and 10 min after the initiation of transcription through addition of T7 RNAP, showing the aggregation of RNA particles. Scale bar: 5 µm. **b** Number and brightness analysis demonstrating aggregation of RNA over time and recruitment of nascent RNA molecules by already existing gel condensates. The brightness per molecule increases for similar intensities as the transcription proceeds after activation, indicating RNA oligomerization (such that oligomers of RNA now move together as opposed to monomers in the beginning of transcription). Number density maps and corresponding to the brightness maps show a higher number density in the middle of the patterns, indicating that these structures grow outwards from nucleation points.

To test whether RNA phase separation arose solely from increased concentration, we compared transcription of two distinct RNA sequences (Figure S2a). Although only one sequence, containing aptameric repeats downstream of the coding region, aggregated spontaneously in transcription buffer at high concentration, both sequences readily phase separated during MBT, demonstrating that confinement within the DNA network facilitates condensation. Further experiments revealed that stable RNA phase separation requires electrostatic crosslinking by multivalent cations, namely magnesium and spermidine. When cation availability within the DNA layer was reduced, RNA transcription proceeded efficiently, but RNA condensates were transient and ultimately escaped the DNA phase at the membrane surface (Figure S2b-c).

Together, these results indicate that RNA undergoes co-transcriptional liquid-to-solid phase exclusion within the DNA network. As nascent RNA oligomerizes and gels, it becomes impenetrable to double-stranded DNA, driving demixing between the two nucleic-acid phases.

### Lipid membranes modulate mechanical feedback between RNA and DNA phases

Lipid membranes are known to display intrinsic affinity for nucleic acids and, in some contexts, to exert catalytic effects on their assembly and reactivity. (8-14) We therefore examined how membrane properties influence RNA patterning within the MBT system. First, we compared POPC (1-palmitoyl-2-oleoyl-sn-glycero-3-phosphocholine) and DPPC (1,2-dipalmitoyl-sn-glycero-3-phosphocholine) membranes, as these lipids exhibit increasing DNA and RNA binding affinity in parallel with increasing transition temperature (Tm). (10, 11, 13) Reducing membrane fluidity increased DNA binding, consistent with previous findings. DNA binding to DPPC membranes was sufficiently strong that cholesterol anchoring was not required. Regardless of membrane fluidity, meandering RNA patterns formed, but the penetration depth of the RNA gel into the DNA layer varied, appearing as reduced anti-localization of the nucleic acids at lower membrane fluidity (Figure 4a).

**Figure 4.**
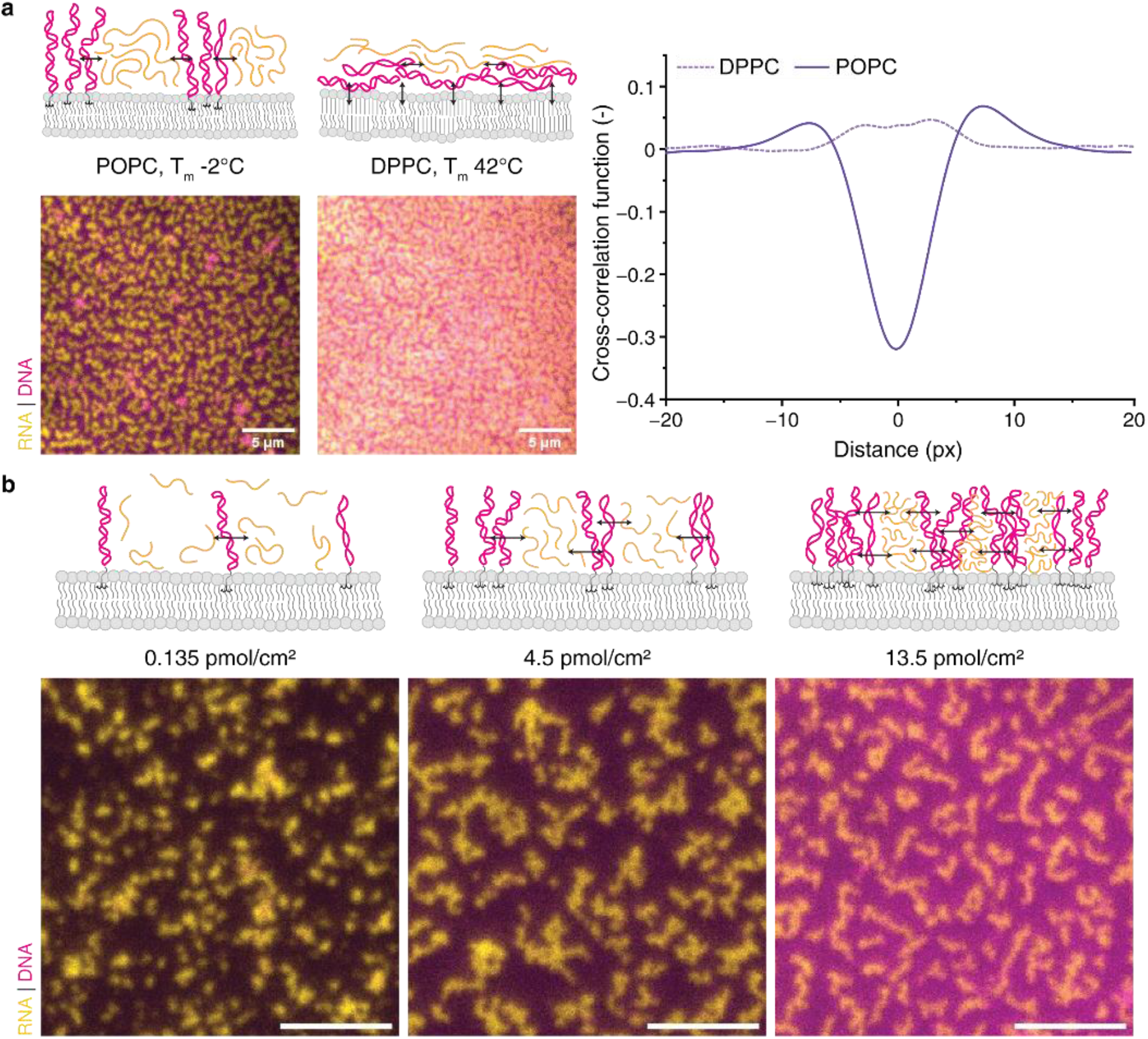
Lipid membranes modulate the feedback between DNA and RNA phases. **a** RNA phase separation on liquid-disordered (POPC, T_m_ = –2 °C) and ripple phase (DPPC, T_m_ = 42 °C) membranes. Cross-correlation analysis shows anti-localization between the DNA and RNA phases reduce with a reduction in the fluidity of the membrane environment. **b** RNA phase separation as a function of DNA density. As the density of tethered DNA at the membrane surface is increased, the morphology of the resulting RNA phases changes. Scale bars: 5 µm.

Membrane headgroup charge exerted a strong effect: RNA patterning occurred exclusively on zwitterionic membranes (POPC), whereas positively charged membranes (DOTAP, 1,2-dioleoyl-3-trimethylammonium-propane) failed to concentrate RNA, and negatively charged membranes (POPG, 1-palmitoyl-2-oleoyl-sn-glycero-3-phospho-(1′-rac-glycerol)) produced homogeneous RNA distributions (Figure S3). We explain this counterintuitive result by cation partitioning. Negative PG headgroups up-concentrate cations at the membrane surface and attract nucleic acids through ionic bridging, whereas positive TAP headgroups likely repel RNA that is passivated by cations in solution.

We next investigated how direct changes in DNA network density affect RNA transcription and patterning on fluid, neutral POPC membranes (Figure 4b). We observed RNA condensation across the full tested range of DNA surface densities (0.135 to 13.5 pmol per cm^2^ of SLB area). In contrast, RNA-phase patterning was more robust at higher surface densities, with 4.5 pmol/cm^2^ and above, leading to more compact and elongated RNA structures. This coincided with an increased initial rate of transcription and faster saturation of denser DNA layers with nascent RNA molecules (Figure S4).

To verify whether the observed effects were limited to single-component membranes, we assembled the MBT system on membranes derived from natural, compositionally complex lipid extracts. For this, we used total lipid extracts from E. coli, soy, and liver to form SLBs (Figure 5a). Each extract is significantly different in lipid composition (main lipids of E. coli: PE, PG, and cardiolipin; soy: PC, PE, PI, and PA; liver: PC, PE, PI, and cholesterol), and containing between up to 20% of unidentified lipids. Strikingly, each membrane produced RNA patterns of distinct morphology: liver extract yielded an “archipelago” pattern similar to POPC membranes, soy extract produced a “spots” pattern, and E. coli extract gave a phase-dependent morphology with predominantly “inverted archipelago” patterns. In addition, we performed MBT on simplified, phase-separated membranes (70% POPE, 25% POPG, 5% cardiolipin) designed to mimic E. coli lipid extract (Figure 5b). Here too, we observed a transition from “inverted archipelago” morphology in the cardiolipin-rich phase to “archipelago” morphology in the cardiolipin-poor phase upon local changes in lipid phase. Notably, at later stages of system evolution, the lipid membrane phase drives even further, leading to almost full exclusion of RNA and enrichment of DNA within the cardiolipin-rich lipid phase.

**Figure 5.**
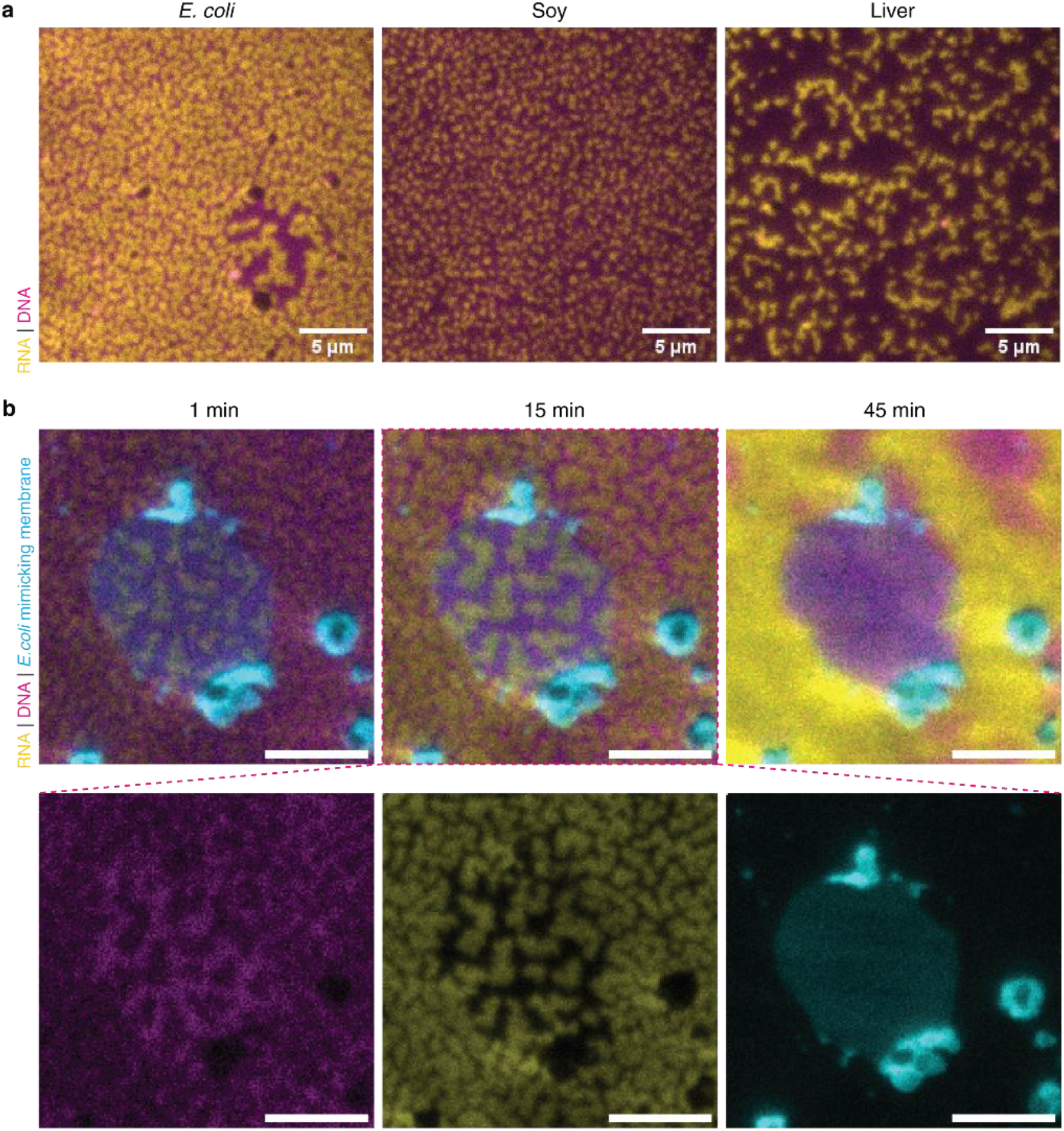
Pattern behavior on various membranes. **a** RNA/ DNA patterns on membranes formed with *E. coli*, soy, and liver lipid extracts. The liver membrane yielded an “archipelago” pattern, the soy membrane produced a “spots” pattern, and the *E. coli* membrane gave a phase-dependent morphology with predominantly “inverted archipelago” patterns. **b** RNA phase separation on *E. coli*-mimicking phase-separating membranes of a simplified lipid composition (70% POPE, 25% POPG, 5% cardiolipin). The morphology of RNA condensates is locally modulated by the underlying membrane phase (the cardiolipin-rich phase forms an island in the middle of the image). Scale bars: 5 µm.

These results highlight the importance of subtle environmental, i.e., electrostatic and mechanical, effects mediated by the membrane. Robust RNA pattern formation requires a DNA network that is both sufficiently fluid to rearrange and sufficiently viscous to resist expansion of the RNA gel. We therefore consider the observed patterning to arise from dynamic interplay of the forces generated by RNA growth and the rearrangement of the surrounding DNA phase. In this picture, the membrane acts not merely as a passive support, but as an active interfacial regulator that modulates DNA-layer fluidity, local cation availability, and the capacity of the surface to concentrate condensed RNA. These properties strongly influence the mechanical feedback that underlies pattern formation and provide a direct explanation for the sensitivity of pattern morphology to membrane composition.

### RNA–protein interactions modulate RNA gelation and sequestration

This raised a further question: what happens to T7 RNA polymerase during the observed nucleic-acid transformation? We addressed this using two experimental approaches: polymerase targeting through promoter placement, and direct localization monitoring using fluorescently labeled polymerase.

First, we compared PCR products in which the transcription start site was placed either close to the membrane anchor or at the opposite end of the template (Figure 6a). In this way, we could force the polymerase to start RNA synthesis in proximity to the membrane and continue outward, or conversely to start farther away and continue toward the membrane. We found that robust patterning occurred when nascent RNA was generated deep within the DNA layer near the membrane. When transcription initiated farther from the membrane, RNA still condensed but escaped the DNA phase more readily. Here, we consider two complementary mechanistic factors. First, electrostatic attraction to the membrane surface may create a short-range energy landscape that traps nascent RNA more efficiently. Second, expansion of the trapped RNA phase may force tethered DNA molecules to move laterally together with their lipid anchors. Alternatively, lateral rearrangement could in principle arise from polymerase action itself. We therefore turned to the second approach and directly monitored polymerase localization during transcription.

**Figure 6.**
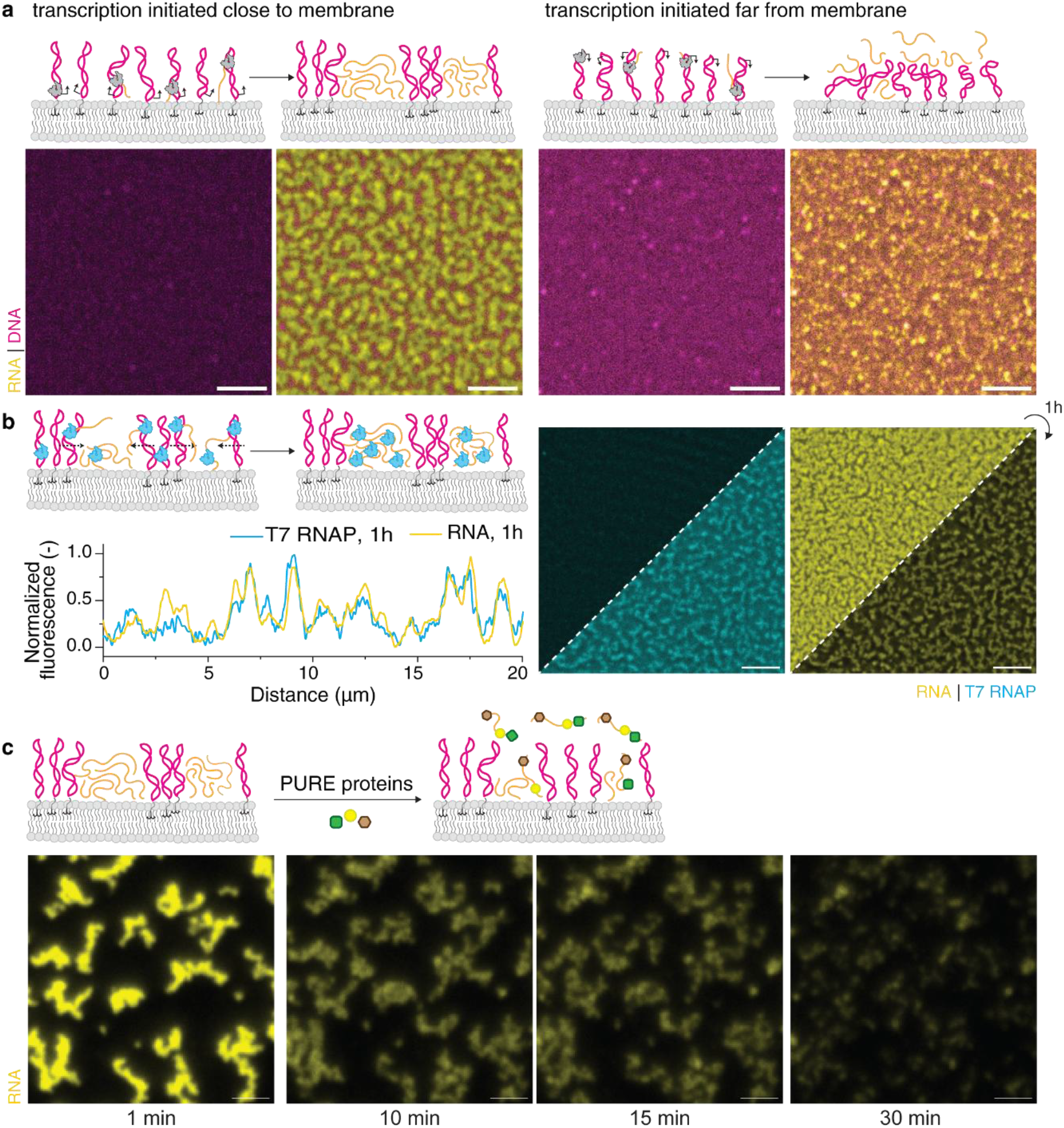
RNA–protein interactions regulate RNA gelation and sequestration. **a** TIRF images, recorded before and after one hour of transcription, demonstrating the phase separation of RNA dependent on targeting of T7 RNA polymerase. The promoter sequence, when placed near the membrane tether (left) or further from the membrane (right), resulted in varying degrees of RNA gelation. Scale bars: 5 µm. **b** Monitoring of T7 RNA polymerase localization during MBT using a fluorescently labelled T7 RNA polymerase (Alexa Fluor 488, cyan). TIRF images were recorded after 5, 15 and 60 min after addition the labelled polymerase. Scale bars: 5 µm. **c** Dissolution of RNA gel phase by PURE proteins. TIRF images were recorded 1, 10, 15 and 30 min after the addition of the PURE protein mixture to a pre-developed RNA pattern. Scale bar: 2 µm.

For this, we used AF488-labeled T7 RNA polymerase. Tracking T7 RNA polymerase revealed its progressive sequestration into the RNA gel. Upon completion of transcription, the polymerase was almost entirely excluded from the DNA phase (Figure 6b). This observation is consistent with the idea that the gel phase is enriched in stable intra- and intermolecular RNA secondary structures arising from RNA-RNA base-pairing and further stabilized by cationic crosslinking, thereby facilitating RNA condensation and DNA-RNA phase separation. At the same time, we also consider the possibility that sequestration is driven, at least in part, by the macroscopic material properties of the condensed RNA phase rather than exclusively by RNA secondary structure.

This led us to the final question: how do proteins engaged in gene expression and regulation interact with the condensed RNA patterns? To test this, after pattern formation we added the PURE-system protein components, comprising a minimal set of enzymes and transcription and translation factors necessary for cell-free expression (22). Strikingly, addition of PURE proteins reversed RNA gelation and restored DNA fluidity (Figure 6c). On one hand, this supported our interpretation that non-specific RNA-RNA interactions contribute substantially to phase separation. On the other, it showed that disruption of those interactions by RNA-binding proteins can be equally robust.

These results suggest that, in the absence of regulatory factors, transcription in a dense DNA network intrinsically drives RNA gelation and can lead to non-specific protein sequestration. In that light, RNA-binding factors may serve a dual function: enabling downstream gene expression while simultaneously preventing spontaneous RNA condensation.

## Conclusions

We introduce a membrane-bound transcription system that serves as a programmable and minimal platform for studying nucleic-acid interactions and emergent behavior at membrane surfaces. Using this platform, we find that RNA phase separation can arise as an intrinsic consequence of transcription under confinement. At the same time, our results argue that the membrane is not merely a passive boundary condition. Rather, the interface actively shapes the onset, morphology, and evolution of DNA-RNA unmixing, likely by modulating nucleic-acid adsorption, lateral mobility, local electrostatics, and cation partitioning. In this view, the membrane acts as an interfacial regulator of nucleic-acid reorganization: it likely biases RNA-RNA interactions, promotes retention of nascent RNA on the membrane surface, and thereby alters the mechanical feedback between the expanding RNA phase and the surrounding DNA network. The phase-dependent switching observed in Figure 5b provides especially direct support for this interpretation, showing that local lipid composition can redirect the patterning pathway and its time evolution.

These findings support a broader biological relevance for the phenomenon. Nuclear and chromatin-associated environments contain multivalent cations and other electrostatic regulators that can favor nucleic-acid compaction and intermolecular association. (23, 24) Our results therefore raise the possibility that transcription in vivo may operate close to a physical instability threshold, at which newly synthesized RNA would tend to condense unless buffered by RNA-binding proteins, RNA processing pathways, and downstream gene-expression machinery. In our minimal system, multivalent cations stabilize condensation, whereas the addition of PURE proteins reverses gelation. Taken together, this suggests that one evolutionary role of RNA-binding and translation-associated factors may be to limit spatial RNA accumulation, thereby helping preserve genome organization while maintaining productive information flow. (23, 25)

Beyond modern cells, the results also resonate with scenarios for the origins and early evolution of life. Surface effects and RNA-lipid interactions are increasingly recognized as important features of prebiotic chemistry, because interfaces can enrich reactants, alter local electrostatic conditions, and create spatially heterogeneous reaction environments. (9) Our findings extend this idea from molecular assembly to dynamic nucleic-acid phase behavior. Surface charge, lipid composition, and membrane phase state can all regulate where RNA accumulates and whether the resulting structures remain dynamic or become arrested, likely by influencing the extent of RNA self-association. Primitive surfaces may therefore have provided not only compartments, but also spatial, temporal, and mechanistic control over RNA organization before the emergence of elaborate protein-based regulation.

More broadly, this work shows how surprisingly complex behavior can emerge from minimal components: DNA, RNA, lipids, and basic electrostatics. The balance of forces underlying the process is experimentally controllable, offering new opportunities for synthetic biology, biophysics, and the bottom-up design of transcriptionally active soft-matter systems. Ultimately, our findings support a simple but far-reaching principle: transcription is a physically active process that produces a polymer prone to phase separation. In confined and crowded environments, continuous RNA synthesis may create a materials-level instability that can drive gelation, phase exclusion, nucleic-acid reorganization, and protein sequestration unless counteracted. We therefore propose that an important function of RNA-binding proteins, RNA processing, and translation machinery may be not only to regulate information flow, but also to buffer against transcription-driven physical collapse or compaction of genome organization. In this view, the molecular complexity of gene expression reflects not only the need to control genetic information, but also the need to maintain physical homeostasis in an inherently self-organizing nucleic-acid system.

## Supporting information

Supplementary figures, DNA sequences, experimental methods

## Acknowledgements

This study was supported by Alexander von Humbold Foundation and European Union under the Marie Skłodowska-Curie Actions Postdoctoral Fellowship (101208282). We thank Beatrix Scheffer and Sandra Ortmeier for help in preparation of DNA templates, lipid mixes and supported lipid bilayers. We thank the MPIB Bioorganic Chemistry and biophysics core facility for performing MS analysis and access to HPLC during chemical synthesis.

## Author information

## Author contributions

A.M. conceived and supervised the project, developed the MBT system, performed or was involved in all described experiments, and drafted and revised the manuscript. T.A.N. performed experiments, analyzed TIRF imaging and FRAP data, and edited the manuscript. Y.Q. performed experiments and analyzed N&B data. S.G. performed experiments and analyzed AFM data. S.B. optimized MBT system assembly and carried out membrane screening experiments. V.B. prepared cell-free expression proteins. J.-H.K. developed scripts for TIRF data analysis. N.K. prepared fluorescently labeled T7 RNA polymerase. P.S. revised the manuscript and provided overall guidance.

## Use of A.I. tools

Large language models (i.e. ChatGPT and Perplexity) were used to assist with literature searches and language editing of human-written draft. The authors take full responsibility for the content and conclusions of this work.

