## Supplementary figures, DNA sequences, experimental methods for "Co-transcriptional Phase Separation of Nucleic Acids at Membrane Surfaces"

T7 RNAP Alexa Fluor 488 maleimide ........ 14

#### Supplementary Figures

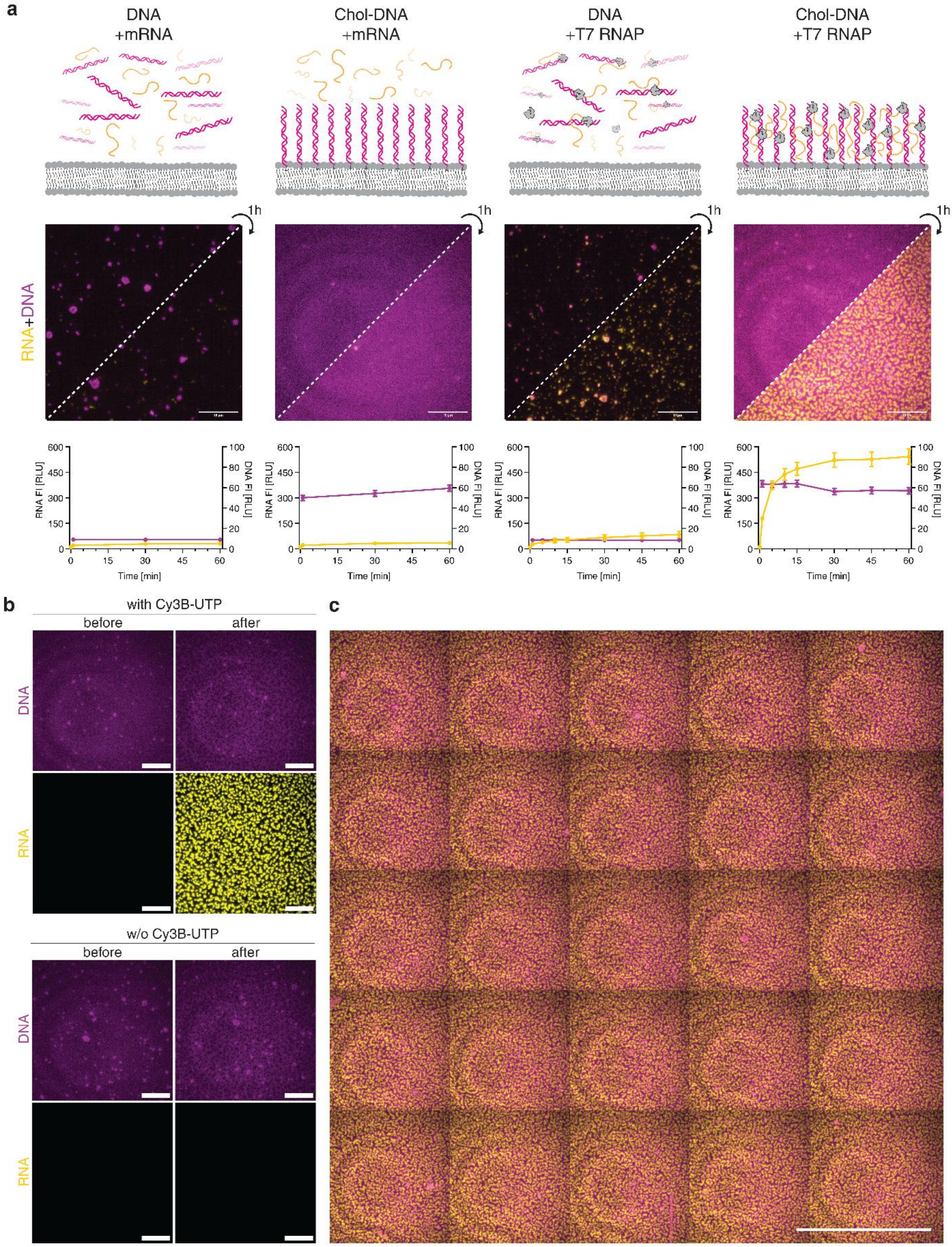

**Figure S1. a** Comparison of MBT with static mixtures and transcription in solution. From left to right: pre-synthesized mRNA was mixed with unmodified DNA and incubated over supported lipid bilayers (SLBs); pre-synthesized mRNA was incubated over SLBs with membrane-tethered DNA; mRNA was transcribed from non-tethered DNA over SLBs; MBT set-up. TIRF images show the systems at the onset of and one hour after incubation/transcription of the systems. Graphs below show fluorescence intensity changes over time for RNA and DNA channels, averaged over 25 independent spots within a sample. Scale bars: 10 µm. **b** MBT with and without labelled nucleotide (Cy3B-UTP)/ RNA. TIRF images show DNA (magenta) and RNA (yellow, if labelled) at the onset of and after one hour of transcription. Scale bars: 10 µm. **c** Tilescan TIRF image of MBT system. Scale bar: 50 µm.

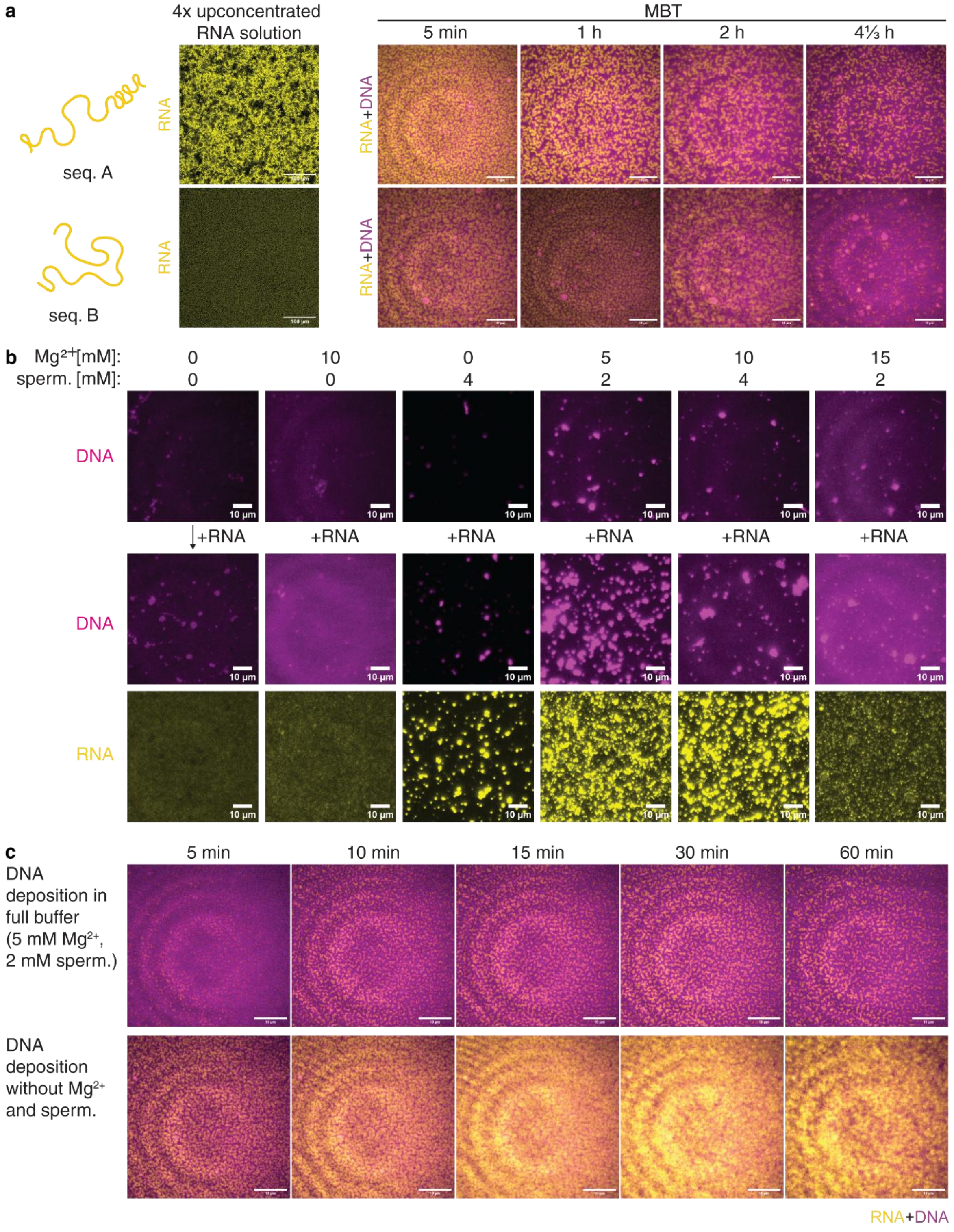

**Figure S2. a** Left - Confocal fluorescence microscopy image of up-concentrated RNA sequences A and B. Right – Transcription of sequences A and B in MBT setting, and sequence dependent temporal evolution and decomposition of RNA patterns. **b** Static system imaging in presence of different concentration of magnesium and spermidine. **c** Comparison of system evolution in dependence on saturation of DNA layer with multivalent cations. To that end, DNA was deposited and pre incubated on top of the POPC lipid membranes in presence or absence of magnesium acetate and spermidine. The full buffer composition was restored shortly before initiation of transcription. Scale bars: 10 µm.

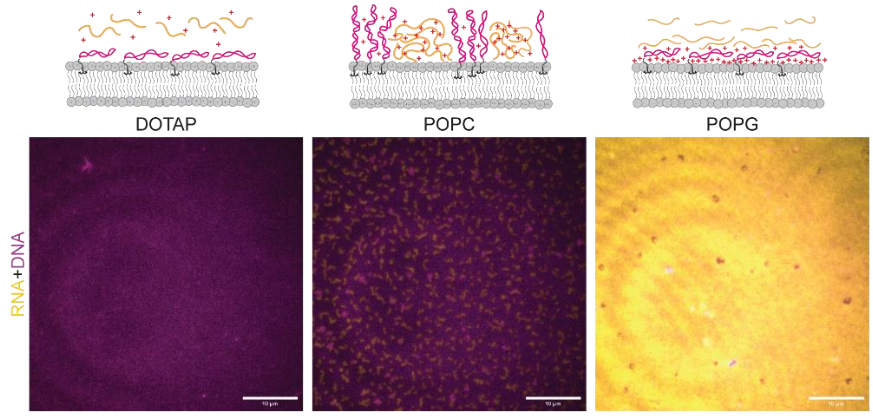

**Figure S3.** TIRF images recorded after 15 min from MTB initiation, demonstrating influence of lipid headgroup charge on patterning and localization of nucleic acids. Scale bars: 10 µm.

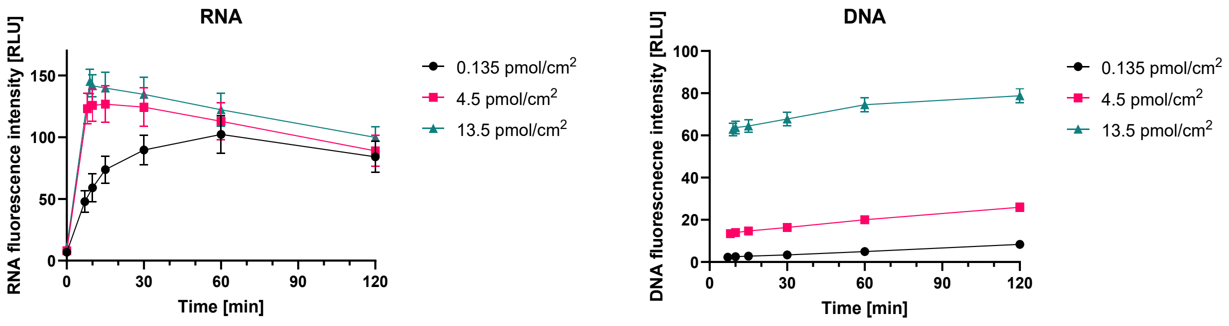

**Figure S4**. Transcription progress in dependence on DNA surface density.

### Plasmid maps and sequences

**pCoofy_msfGFP** (code: PL_B)

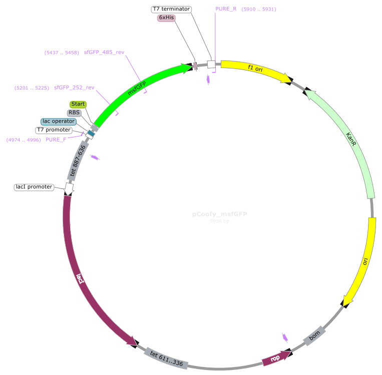

tggcgaatgggacgcgccctgtagcggcgcattaagcgcggcgggtgtggtggttacgcgcagcgtgaccgctacacttgcc
agcgccctagcgcccgctcctttcgctttcttcccttcctttctcgccacgttcgccggctttccccgtcaagctctaaatcgggg
ctccctttagggttccgatttagtgctttacggcacctcgaccccaaaaaacttgattagggtgatggttcacgtagtgggccatcg
ccctgatagacggtttttcgccctttgacgttggagtccacgttctttaatagtggactcttgttccaaactggaacaacactcaacc
ctatctcggtctattcttttgatttataagggattttgccgatttcggcctattggttaaaaaatgagctgatttaacaaaaatttaacgcg
aattttaacaaaatattaacgtttacaatttcaggtggcacttttcggggaaatgtgcgcggaacccctatttgtttatttttctaaataca
ttcaaatatgtatccgctcatgaattaattcttagaaaaactcatcgagcatcaaatgaaactgcaatttattcatatcaggattatcaa
taccatatttttgaaaaagccgtttctgtaatgaaggagaaaactcaccgaggcagttccataggatggcaagatcctggtatcgg
tctgcgattccgactcgtccaacatcaatacaacctattaatttcccctcgtcaaaaataaggttatcaagtgagaaatcaccatga
gtgacgactgaatccggtgagaatggcaaaagtttatgcatttctttccagacttgttcaacaggccagccattacgctcgtcatca
aaatcactcgcatcaaccaaaccgttattcattcgtgattgcgcctgagcgagacgaaatacgcgatcgctgttaaaaggacaat
tacaaacaggaatcgaatgcaaccggcgcaggaacactgccagcgcatcaacaatattttcacctgaatcaggatattcttctaa
tacctggaatgctgttttcccggggatcgcagtggtgagtaaccatgcatcatcaggagtacggataaaatgcttgatggtcggaa
gaggcataaattccgtcagccagtttagtctgaccatctcatctgtaacatcattggcaacgctacctttgccatgtttcagaaacaa
ctctggcgcatcgggcttcccatacaatcgatagattgtcgcacctgattgcccgacattatcgcgagcccatttatacccatataa
atcagcatccatgttggaatttaatcgcggcctagagcaagacgtttcccgttgaatatggctcataacaccccttgtattactgttta
tgtaagcagacagttttattgttcatgaccaaaatcccttaacgtgagttttcgttccactgagcgtcagaccccgtagaaaagatca
aaggatcttcttgagatcctttttttctgcgcgtaatctgctgcttgcaaacaaaaaaaccaccgctaccagcggtggtttgtttgccg
gatcaagagctaccaactctttttccgaaggtaactggcttcagcagagcgcagataccaaatactgtccttctagtgtagccgta
gttaggccaccacttcaagaactctgtagcaccgcctacatacctcgctctgctaatcctgttaccagtggctgctgccagtggcg
ataagtcgtgtcttaccgggttggactcaagacgatagttaccggataaggcgcagcggtcgggctgaacggggggttcgtgc
acacagcccagcttggagcgaacgacctacaccgaactgagatacctacagcgtgagctatgagaaagcgccacgcttccc
gaagggagaaaggcggacaggtatccggtaagcggcagggtcggaacaggagagcgcacgagggagcttccaggggga
aacgcctggtatctttatagtcctgtcgggtttcgccacctctgacttgagcgtcgatttttgtgatgctcgtcaggggggcggagcc
tatggaaaaacgccagcaacgcggcctttttacggttcctggccttttgctggccttttgctcacatgttctttcctgcgttatcccctg
attctgtggataaccgtattaccgcctttgagtgagctgataccgctcgccgcagccgaacgaccgagcgcagcgagtcagtg
agcgaggaagcggaagagcgcctgatgcggtattttctccttacgcatctgtgcggtatttcacaccgcatatatggtgcactctc
agtacaatctgctctgatgccgcatagttaagccagtatacactccgctatcgctacgtgactgggtcatggctgcgccccgaca
cccgccaacacccgctgacgcgccctgacgggcttgtctgctcccggcatccgcttacagacaagctgtgaccgtctccggg
agctgcatgtgtcagaggttttcaccgtcatcaccgaaacgcgcgaggcagctgcggtaaagctcatcagcgtggtcgtgaag
cgattcacagatgtctgcctgttcatccgcgtccagctcgttgagtttctccagaagcgttaatgtctggcttctgataaagcgggcc
atgttaagggcggttttttcctgtttggtcactgatgcctccgtgtaagggggatttctgttcatgggggtaatgataccgatgaaacg
agagaggatgctcacgatacgggttactgatgatgaacatgcccggttactggaacgttgtgagggtaaacaactggcggtatg
gatgcggcgggaccagagaaaaatcactcagggtcaatgccagcgcttcgttaatacagatgtaggtgttccacagggtagcc
agcagcatcctgcgatgcagatccggaacataatggtgcagggcgctgacttccgcgtttccagactttacgaaacacggaaa
ccgaagaccattcatgttgttgctcaggtcgcagacgttttgcagcagcagtcgcttcacgttcgctcgcgtatcggtgattcattct
gctaaccagtaaggcaaccccgccagcctagccgggtcctcaacgacaggagcacgatcatgcgcacccgtggggccgcc

atgccggcgataatggcctgcttctcgccgaaacgtttggtggcgggaccagtgacgaaggcttgagcgagggcgtgcaaga
ttccgaataccgcaagcgacaggccgatcatcgtcgcgctccagcgaaagcggtcctcgccgaaaatgacccagagcgctg
ccggcacctgtcctacgagttgcatgataaagaagacagtcataagtgcggcgacgatagtcatgccccgcgcccaccggaa
ggagctgactgggttgaaggctctcaagggcatcggtcgagatcccggtgcctaatgagtgagctaacttacattaattgcgttg
cgctcactgcccgctttccagtcgggaaacctgtcgtgccagctgcattaatgaatcggccaacgcgcggggagaggcggttt
gcgtattgggcgccagggtggtttttcttttcaccagtgagacgggcaacagctgattgcccttcaccgcctggccctgagagag
ttgcagcaagcggtccacgctggtttgccccagcaggcgaaaatcctgtttgatggtggttaacggcgggatataacatgagctg
tcttcggtatcgtcgtatcccactaccgagatatccgcaccaacgcgcagcccggactcggtaatggcgcgcattgcgcccag
cgccatctgatcgttggcaaccagcatcgcagtgggaacgatgccctcattcagcatttgcatggtttgttgaaaaccggacatg
gcactccagtcgccttcccgttccgctatcggctgaatttgattgcgagtgagatatttatgccagccagccagacgcagacgcg
ccgagacagaacttaatgggcccgctaacagcgcgatttgctggtgacccaatgcgaccagatgctccacgcccagtcgcgt
accgtcttcatgggagaaaataatactgttgatgggtgtctggtcagagacatcaagaaataacgccggaacattagtgcaggc
agcttccacagcaatggcatcctggtcatccagcggatagttaatgatcagcccactgacgcgttgcgcgagaagattgtgcac
cgccgctttacaggcttcgacgccgcttcgttctaccatcgacaccaccacgctggcacccagttgatcggcgcgagatttaatc
gccgcgacaatttgcgacggcgcgtgcagggccagactggaggtggcaacgccaatcagcaacgactgtttgcccgccagt
tgttgtgccacgcggttgggaatgtaattcagctccgccatcgccgcttccactttttcccgcgttttcgcagaaacgtggctggcct
ggttcaccacgcgggaaacggtctgataagagacaccggcatactctgcgacatcgtataacgttactggtttcacattcaccac
cctgaattgactctcttccgggcgctatcatgccataccgcgaaaggttttgcgccattcgatggtgtccgggatctcgacgctctc
ccttatgcgactcctgcattaggaagcagcccagtagtaggttgaggccgttgagcaccgccgccgcaaggaatggtgcatgc
aaggagatggcgcccaacagtcccccggccacggggcctgccaccatacccacgccgaaacaagcgctcatgagcccga
agtggcgagcccgatcttccccatcggtgatgtcggcgatataggcgccagcaaccgcacctgtggcgccggtgatgccggc
cacgatgcgtccggcgtagaggatcgagatctcgatcccgcgaaattaatacgactcactataggggaattgtgagcggataa
caattcccctctagaaataattttgtttaactttaagaaggagatataccatgGTTAGCAAAGGCGAAGAACTGTTTA
CCGGTGTTGTTCCGATTCTGGTTGAACTGGATGGTGATGTTAATGGCCACAAATTTTCAGT
TCGTGGTGAAGGCGAAGGTGATGCAACCAATGGTAAACTGACCCTGAAATTTATCTGTAC
CACCGGCAAACTGCCGGTTCCGTGGCCGACACTGGTTACCACACTGACCTATGGTGTTC
AGTGTTTTAGCCGTTATCCGGATCATATGAAACGCCACGATTTTTTCAAAAGCGCAATGCC
GGAAGGTTATGTTCAAGAACGTACCATCTCCTTTAAAGATGATGGCACCTATAAAACCCGT
GCCGAAGTTAAATTTGAAGGTGATACCCTGGTGAATCGCATTGAACTGAAAGGCATCGAT
TTCAAAGAAGATGGTAATATCCTGGGCCACAAACTGGAATATAATTTCAATAGCCACAACG
TGTATATCACCGCAGACAAACAGAAAAATGGCATCAAAGCCAACTTTAAGATTCGGCATA
ATGTTGAAGATGGCAGCGTTCAGCTGGCAGATCATTATCAGCAGAATACCCCGATTGGTG
ATGGTCCGGTTCTGCTGCCGGATAATCATTATCTGAGCACCCAGAGCAAACTGAGCAAAG
ATCCGAATGAAAAACGTGATCACATGGTGCTGCTGGAATTTGTTACCGCAGCAGGTATTA
CCCATGGTATGGATGAACTGTATAAAtgagcactcgagcaccaccaccaccaccactgagatccggctgcta
acaaagcccgaaaggaagctgagttggctgctgccaccgctgagcaataactagcataaccccttggggcctctaaacgggt
cttgaggggttttttgctgaaaggaggaactatatccggat

**pPT1_ NeonCyan_link_F30st_MangoA10U_F30mid_MangoA10U_F30end_link** (code: PL_D)

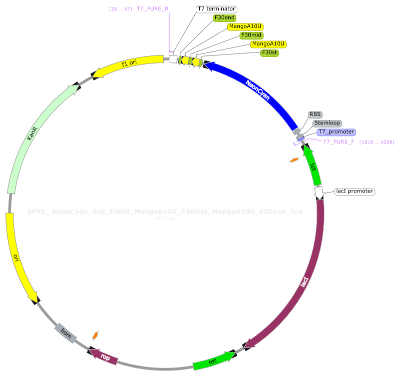

atccggatatagttcctcctttcagcaaaaaacccctcaagacccgtttagaggccccaaggggttatgctagttagcagccgTT
GCCATGAATGATCCGGCACGTACGAATATACCACATACCAAACCTTCCTTCGTACGTGCC
GGATCATCAGAGTATGTGGGGGCACGTACGAATATACCACATACCAAACCTTCCTTCGTA
CGTGCCCCCACATACACATGGCAAgatcaccctTTACTTGTATAACTCGTCCATACCCATCAC
ATCCGTAAAAGCCTTCTGCCATTCTTTAAAATTTAACTCTGTCTCGCTATGTTTCAGCTCTG
TCTTGCGGAACACGTAGACTGGCTGGTTCTTTAAATAGTTAGCTGCCATGGGTTTAGCGA
AGGTATATGTTGTACGGGCCGTGCTGCGGTAACGCTCGCCATTGCCGGTCGTGTAGGAC
CACTTAAAGGTGCTGATGATGGTTTTGTCGTTAGGATATGTCTTTTTAGACCAGCAAGAGT
CTACCGCAGTCAGGCTATTGGTCATCACAGGACCATCGGCTGGGAATCCTGTGCCTTTAA
CCTGGGCTTCCCCTTTGATGTGCGACCCTTCATAAGTATAGCGATAGTTCACTGTTAAAGA
CGCACCGTCCTCAAACTGCATAGTGCGATGCACTTGATATCCGCTGCCGTCTACCATGGC
AGCCTGGAAGGGGCTCATTCCATCAGGGTACGGAAGATACTGATGAAAACCCCATCCCA
TATGTGGGACAAGAATCCACGGACTAAACTGCAGATCCCCTTTCGTGCTCTTCAGATTAA
GTTCCTCATATCCTTCGTTAGGGTTACCCGTACCTTGCCCTACCATATCAAAGTCTACCCC
GTTAATGCTACCGAAAATATGTAATTCATGAGTTGCAGGTAATGATGCCATATTATCTTCTT
CACCTTTTGAAACCATggtatatctccttcttaaagttaaacaaaattatttctagagggAAACCGTTGTGGTCTc
cctatagtgagtcgtattaatttcgcgggatcgagatctcgatcctctacgccggacgcatcgtggccggcatcaccggcgcca
caggtgcggttgctggcgcctatatcgccgacatcaccgatggggaagatcgggctcgccacttcgggctcatgagcgcttgtt
tcggcgtgggtatggtggcaggccccgtggccgggggactgttgggcgccatctccttgcatgcaccattccttgcggcggcg
gtgctcaacggcctcaacctactactgggctgcttcctaatgcaggagtcgcataagggagagcgtcgagatcccggacacca
tcgaatggcgcaaaacctttcgcggtatggcatgatagcgcccggaagagagtcaattcagggtggtgaatgtgaaaccagta
acgttatacgatgtcgcagagtatgccggtgtctcttatcagaccgtttcccgcgtggtgaaccaggccagccacgtttctgcgaa
aacgcgggaaaaagtggaagcggcgatggcggagctgaattacattcccaaccgcgtggcacaacaactggcgggcaaac
agtcgttgctgattggcgttgccacctccagtctggccctgcacgcgccgtcgcaaattgtcgcggcgattaaatctcgcgccga
tcaactgggtgccagcgtggtggtgtcgatggtagaacgaagcggcgtcgaagcctgtaaagcggcggtgcacaatcttctcg
cgcaacgcgtcagtgggctgatcattaactatccgctggatgaccaggatgccattgctgtggaagctgcctgcactaatgttcc
ggcgttatttcttgatgtctctgaccagacacccatcaacagtattattttctcccatgaagacggtacgcgactgggcgtggagca
tctggtcgcattgggtcaccagcaaatcgcgctgttagcgggcccattaagttctgtctcggcgcgtctgcgtctggctggctggc
ataaatatctcactcgcaatcaaattcagccgatagcggaacgggaaggcgactggagtgccatgtccggttttcaacaaacca
tgcaaatgctgaatgagggcatcgttcccactgcgatgctggttgccaacgatcagatggcgctgggcgcaatgcgcgccatta
ccgagtccgggctgcgcgttggtgcggatatctcggtagtgggatacgacgataccgaagacagctcatgttatatcccgccgtt
aaccaccatcaaacaggattttcgcctgctggggcaaaccagcgtggaccgcttgctgcaactctctcagggccaggcggtga
agggcaatcagctgttgcccgtctcactggtgaaaagaaaaaccaccctggcgcccaatacgcaaaccgcctctccccgcgc
gttggccgattcattaatgcagctggcacgacaggtttcccgactggaaagcgggcagtgagcgcaacgcaattaatgtaagtt
agctcactcattaggcaccgggatctcgaccgatgcccttgagagccttcaacccagtcagctccttccggtgggcgcggggc
atgactatcgtcgccgcacttatgactgtcttctttatcatgcaactcgtaggacaggtgccggcagcgctctgggtcattttcggcg
aggaccgctttcgctggagcgcgacgatgatcggcctgtcgcttgcggtattcggaatcttgcacgccctcgctcaagccttcgt
cactggtcccgccaccaaacgtttcggcgagaagcaggccattatcgccggcatggcggccccacgggtgcgcatgatcgtg
ctcctgtcgttgaggacccggctaggctggcggggttgccttactggttagcagaatgaatcaccgatacgcgagcgaacgtga
agcgactgctgctgcaaaacgtctgcgacctgagcaacaacatgaatggtcttcggtttccgtgtttcgtaaagtctggaaacgc
ggaagtcagcgccctgcaccattatgttccggatctgcatcgcaggatgctgctggctaccctgtggaacacctacatctgtatta
acgaagcgctggcattgaccctgagtgatttttctctggtcccgccgcatccataccgccagttgtttaccctcacaacgttccagt
aaccgggcatgttcatcatcagtaacccgtatcgtgagcatcctctctcgtttcatcggtatcattacccccatgaacagaaatccc
ccttacacggaggcatcagtgaccaaacaggaaaaaaccgcccttaacatggcccgctttatcagaagccagacattaacgct
tctggagaaactcaacgagctggacgcggatgaacaggcagacatctgtgaatcgcttcacgaccacgctgatgagctttacc
gcagctgcctcgcgcgtttcggtgatgacggtgaaaacctctgacacatgcagctcccggagacggtcacagcttgtctgtaag
cggatgccgggagcagacaagcccgtcagggcgcgtcagcgggtgttggcgggtgtcggggcgcagccatgacccagtca
cgtagcgatagcggagtgtatactggcttaactatgcggcatcagagcagattgtactgagagtgcaccatatatgcggtgtgaa
ataccgcacagatgcgtaaggagaaaataccgcatcaggcgctcttccgcttcctcgctcactgactcgctgcgctcggtcgttc
ggctgcggcgagcggtatcagctcactcaaaggcggtaatacggttatccacagaatcaggggataacgcaggaaagaacat
gtgagcaaaaggccagcaaaaggccaggaaccgtaaaaaggccgcgttgctggcgtttttccataggctccgcccccctgac
gagcatcacaaaaatcgacgctcaagtcagaggtggcgaaacccgacaggactataaagataccaggcgtttccccctggaa

gctccctcgtgcgctctcctgttccgaccctgccgcttaccggatacctgtccgcctttctcccttcgggaagcgtggcgctttctc
atagctcacgctgtaggtatctcagttcggtgtaggtcgttcgctccaagctgggctgtgtgcacgaaccccccgttcagcccga
ccgctgcgccttatccggtaactatcgtcttgagtccaacccggtaagacacgacttatcgccactggcagcagccactggtaa
caggattagcagagcgaggtatgtaggcggtgctacagagttcttgaagtggtggcctaactacggctacactagaaggacagt
atttggtatctgcgctctgctgaagccagttaccttcggaaaaagagttggtagctcttgatccggcaaacaaaccaccgctggta
gcggtggtttttttgtttgcaagcagcagattacgcgcagaaaaaaaggatctcaagaagatcctttgatcttttctacggggtctga
cgctcagtggaacgaaaactcacgttaagggattttggtcatgaacaataaaactgtctgcttacataaacagtaatacaaggggt
gttatgagccatattcaacgggaaacgtcttgctctaggccgcgattaaattccaacatggatgctgatttatatgggtataaatggg
ctcgcgataatgtcgggcaatcaggtgcgacaatctatcgattgtatgggaagcccgatgcgccagagttgtttctgaaacatgg
caaaggtagcgttgccaatgatgttacagatgagatggtcagactaaactggctgacggaatttatgcctcttccgaccatcaagc
attttatccgtactcctgatgatgcatggttactcaccactgcgatccccgggaaaacagcattccaggtattagaagaatatcctg
attcaggtgaaaatattgttgatgcgctggcagtgttcctgcgccggttgcattcgattcctgtttgtaattgtccttttaacagcgatcg
cgtatttcgtctcgctcaggcgcaatcacgaatgaataacggtttggttgatgcgagtgattttgatgacgagcgtaatggctggcc
tgttgaacaagtctggaaagaaatgcataaacttttgccattctcaccggattcagtcgtcactcatggtgatttctcacttgataacc
ttatttttgacgaggggaaattaataggttgtattgatgttggacgagtcggaatcgcagaccgataccaggatcttgccatcctatg
gaactgcctcggtgagttttctccttcattacagaaacggctttttcaaaaatatggtattgataatcctgatatgaataaattgcagttt
catttgatgctcgatgagtttttctaagaattaattcatgagcggatacatatttgaatgtatttagaaaaataaacaaataggggttcc
gcgcacatttccccgaaaagtgccacctgaaattgtaaacgttaatattttgttaaaattcgcgttaaatttttgttaaatcagctcatttt
ttaaccaataggccgaaatcggcaaaatcccttataaatcaaaagaatagaccgagatagggttgagtgttgttccagtttggaac
aagagtccactattaaagaacgtggactccaacgtcaaagggcgaaaaaccgtctatcagggcgatggcccactacgtgaac
catcaccctaatcaagttttttggggtcgaggtgccgtaaagcactaaatcggaaccctaaagggagcccccgatttagagcttg
acggggaaagccggcgaacgtggcgagaaaggaagggaagaaagcgaaaggagcgggcgctagggcgctggcaagtg
tagcggtcacgctgcgcgtaaccaccacacccgccgcgcttaatgcgccgctacagggcgcgtcccattcgcca

#### List of oligonucleotides

| Code | full name | 5' modification | Tm | sequence |
|---|---|---|---|---|
| B01F | PURE_for | none | 60 | CCCGCGAAATTAATACGACTCAC |
| B01R | PURE_rev | none | 59 | CAAAAAACCCCTCAAGACCCGT |
| B02F | Chol-PURE_for | Cholesteryl TEG | 60 | CCCGCGAAATTAATACGACTCAC |
| B02R | Chol-PURE_rev | Cholesteryl TEG | 59 | CAAAAAACCCCTCAAGACCCGT |
| B03F | A647N-PURE_for | ATTO647N | 60 | CCCGCGAAATTAATACGACTCAC |
| B03R | A647N-PURE_rev | ATTO647N | 59 | CAAAAAACCCCTCAAGACCCGT |
| B05F | N3-PURE_for | Azide | 60 | CCCGCGAAATTAATACGACTCAC |
| B05R | N3-PURE_rev | Azide | 59 | CAAAAAACCCCTCAAGACCCGT |
| D01R | sfGFP_252_rev | none | 59 | CCGGTGGTACAGATAAATTTCAGGG |
| D02R | Chol-sfGFP_252_rev | Cholesteryl TEG | 59 | CCGGTGGTACAGATAAATTTCAGGG |
| D03R | N3-sfGFP_252_rev | Azide | 59 | CCGGTGGTACAGATAAATTTCAGGG |
| D04R | A647N-sfGFP_252_rev | ATTO647N | 59 | CCGGTGGTACAGATAAATTTCAGGG |
| D11R | sfGFP_485_rev | none | 60 | CGATGCCTTTCAGTTCAATGCG |
| D12R | Chol-sfGFP_485_rev | Cholesteryl TEG | 60 | CGATGCCTTTCAGTTCAATGCG |
| D13R | N3-sfGFP_485_rev | Azide | 60 | CGATGCCTTTCAGTTCAATGCG |
| D14R | A647N-sfGFP_485_rev | ATTO647N | 60 | CGATGCCTTTCAGTTCAATGCG |

#### List of DNA templates products and sequences

| Code | primer For | primer Rev | For_mod | Rev_mod | Template | size [bp] | C @ $A_{260}$=1 [nM] |
|---|---|---|---|---|---|---|---|
| B01 | B01F | B01R | no | no | | | |
| B02 | B03F | B01R | A647N | no | | | |
| B03 | B01F | B03R | no | A647N | | | |
| B04 | B01F | B02R | no | Chol | | | |
| B05 | B02F | B01R | Chol | no | PL_B | 955 | 82 |
| B06 | B03F | B02R | A647N | Chol | | | |
| B07 | B02F | B03R | Chol | A647N | | | |
| B08 | B03F | B05R | A647N | N3 | | | |
| B09 | B05F | B03R | N3 | A647N | | | |
| B10 | B01F | D01R | no | no | | | |
| B11 | B03F | D01R | A647N | no | | | |
| B12 | B03F | D02R | A647N | Chol | | | |
| B13 | B03F | D03R | A647N | N3 | PL_B | 252 | 310 |
| B14 | B01F | D04R | no | A647N | | | |
| B15 | B02F | D04R | Chol | A647N | | | |
| B16 | B05F | D04R | N3 | A647N | | | |
| B20 | B01F | D11R | no | no | | | |
| B21 | B03F | D11R | A647N | no | | | |
| B22 | B03F | D12R | A647N | Chol | | | |
| B23 | B03F | D13R | A647N | N3 | PL_B | 485 | 161 |
| B24 | B01F | D14R | no | A647N | | | |
| B25 | B02F | D14R | Chol | A647N | | | |
| B26 | B05F | D14R | N3 | A647N | | | |
| D01 | B01F | B01R | no | no | | | |
| D02 | B01F | B02R | A647N | no | | | |
| D03 | B01F | B03R | no | A647N | | | |
| D04 | B01F | B02R | no | Chol | | | |
| D05 | B02F | B01R | Chol | no | PL_D | 1003 | 78 |
| D06 | B03F | B02R | A647N | Chol | | | |
| D07 | B02F | B03R | Chol | A647N | | | |
| D08 | B03F | B05R | A647N | N3 | | | |
| D09 | B05F | B03R | N3 | A647N | | | |

| Code | Full sequence |
|---|---|
| B01-B09 | cccgcgaaattaatacgactcactataggggaattgtgagcggataacaattcccctctagaaataattttgtttaact<br>ttaagaaggagatataccATGAGTAAAGGAGAAGAATTATTTACTGGAGTTGTCCCAATT<br>CTTGTTGAATTAGATGGTGATGTTAATGGGCACAAATTTTCTGTCCGTGGAGAGG<br>GTGAAGGTGATGCAACAAACGGAAAACTTACCCTTAAATTTATTTGCACTACTG<br>GAAAACTACCTGTTCCATGGCCAACACTTGTCACTACTTTAACTTATGGTGTTCA<br>ATGCTTTTCCCGTTATCCGGATCACATGAAACGGCATGACTTTTTCAAGAGTGC<br>CATGCCCGAAGGTTATGTACAGGAACGCACTATATCTTTCAAAGATGACGGGAC<br>CTACAAGACGCGTGCTGAAGTCAAGTTTGAAGGTGATACCCTTGTTAATCGTAT<br>CGAGTTAAAAGGTATTGATTTTAAAGAAGATGGAAACATTCTCGGACACAAACTC<br>GAGTACAACTTTAACTCACACAATGTATACATCACGGCAGACAAACAAAAGAAT<br>GGAATCAAAGCTAACTTCAAAATTCGCCACAACGTTGAAGATGGATCCGTTCAA<br>CTAGCAGACCATTATCAACAAAATACTCCAATTGGCGATGGCCCTGTCCTTTTA<br>CCAGACAACCATTACCTGTCGACACAATCTAAACTTTCGAAAGATCCCAACGAA<br>AAGCGTGACCACATGGTCCTTCTTGAGTTTGTAACTGCTGCTGGGATTACACAT |

| | |
|---|---|
| | GGCATGGATGAGCTCTACAAAtgagcactcgagcaccaccaccaccaccactgagatccggctgc<br>taacaaagcccgaaaggaagctgagttggctgctgccaccgctgagcaataactagcataaccccttggggcct<br>ctaaacgggtcttgaggggttttttg |
| B11-B16 | cccgcgaaattaatacgactcactataggggaattgtgagcggataacaattcccctctagaaataattttgtttaact<br>ttaagaaggagatataccatgGTTAGCAAAGGCGAAGAACTGTTTACCGGTGTTGTTCCG<br>ATTCTGGTTGAACTGGATGGTGATGTTAATGGCCACAAATTTTCAGTTCGTGGTG<br>AAGGCGAAGGTGATGCAACCAATGGTAAACTGACCCTGAAATTTATCTGTACCA<br>CCGG |
| B20-B26 | cccgcgaaattaatacgactcactataggggaattgtgagcggataacaattcccctctagaaataattttgtttaact<br>ttaagaaggagatataccatgGTTAGCAAAGGCGAAGAACTGTTTACCGGTGTTGTTCCG<br>ATTCTGGTTGAACTGGATGGTGATGTTAATGGCCACAAATTTTCAGTTCGTGGTG<br>AAGGCGAAGGTGATGCAACCAATGGTAAACTGACCCTGAAATTTATCTGTACCA<br>CCGGCAAACTGCCGGTTCCGTGGCCGACACTGGTTACCACACTGACCTATGGT<br>GTTCAGTGTTTTAGCCGTTATCCGGATCATATGAAACGCCACGATTTTTTCAAAA<br>GCGCAATGCCGGAAGGTTATGTTCAAGAACGTACCATCTCCTTTAAAGATGATG<br>GCACCTATAAAACCCGTGCCGAAGTTAAATTTGAAGGTGATACCCTGGTGAATC<br>GCATTGAACTGAAAGGCATCG |
| D01-D09 | caaaaaacccctcaagacccgtttagaggccccaaggggttatgctagttagcagccgTTGCCATGAATG<br>ATCCGGCACGTACGAATATACCACATACCAAACCTTCCTTCGTACGTGCCGGAT<br>CATCAGAGTATGTGGGGGCACGTACGAATATACCACATACCAAACCTTCCTTCG<br>TACGTGCCCCCACATACACATGGCAAgatcaccctTTACTTGTATAACTCGTCCATA<br>CCCATCACATCCGTAAAAGCCTTCTGCCATTCTTTAAAATTTAACTCTGTCTCGC<br>TATGTTTCAGCTCTGTCTTGCGGAACACGTAGACTGGCTGGTTCTTTAAATAGTT<br>AGCTGCCATGGGTTTAGCGAAGGTATATGTTGTACGGGCCGTGCTGCGGTAAC<br>GCTCGCCATTGCCGGTCGTGTAGGACCACTTAAAGGTGCTGATGATGGTTTTGT<br>CGTTAGGATATGTCTTTTTAGACCAGCAAGAGTCTACCGCAGTCAGGCTATTGG<br>TCATCACAGGACCATCGGCTGGGAATCCTGTGCCTTTAACCTGGGCTTCCCCTT<br>TGATGTGCGACCCTTCATAAGTATAGCGATAGTTCACTGTTAAAGACGCACCGT<br>CCTCAAACTGCATAGTGCGATGCACTTGATATCCGCTGCCGTCTACCATGGCAG<br>CCTGGAAGGGGCTCATTCCATCAGGGTACGGAAGATACTGATGAAAACCCCAT<br>CCCATATGTGGGACAAGAATCCACGGACTAAACTGCAGATCCCCTTTCGTGCTC<br>TTCAGATTAAGTTCCTCATATCCTTCGTTAGGGTTACCCGTACCTTGCCCTACCA<br>TATCAAAGTCTACCCCGTTAATGCTACCGAAAATATGTAATTCATGAGTTGCAGG<br>TAATGATGCCATATTATCTTCTTCACCTTTTGAAACCATggtatatctccttcttaaagttaaa<br>caaaattatttctagagggAAACCGTTGTGGTCTccctatagtgagtcgtatta |

#### Cy3B-UTP synthesis

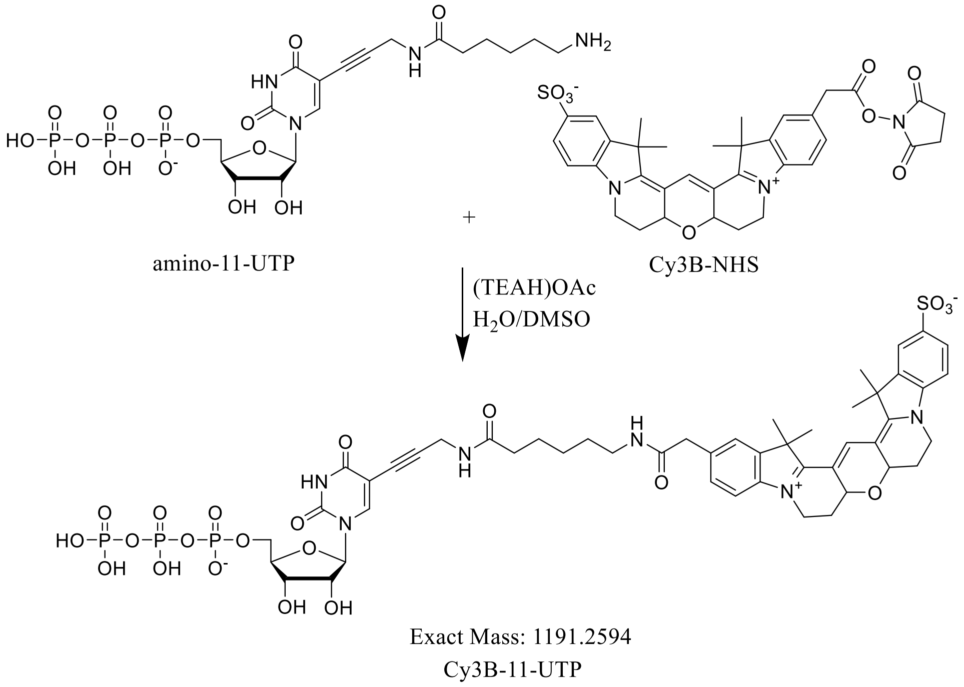

Solutions of amino-11-UTP (19.2 mM, ~2.55 mg / 126 µl of MQ water, $E_{288}$= 12 $mM^{-1}cm^{-1}$, Lumiprobe) and Cy3B-NHS (24.2 mM, ~1.15 mg / 80 µl of anhydrous DMSO, $E_{559}$= 121 $mM^{-1}cm^{-1}$, $E_{290}$~ 9 $mM^{-1}cm^{-1}$ Lumiprobe) were prepared, and their concentration was measured with nanodrop. Two reaction mixtures were prepared by combining amino-11-UTP (5µl, 19.2 mM in water), Cy3B-NHS (5µl, 24.2 mM in DMSO) and triethylammonium acetate (1 µl, 1 M, pH 7.0 or 9.2) solutions (11 µl final volume). The mixtures were vortexed at 100 RPM and 30°C for 1 h and subsequently cooled to room temperature (40 min). Next, the reactions were quenched by dilution with solution of ammonium acetate (90µl, 100 mM, pH 6.7) and resolved using optimized HPLC conditions: column Kinetex 5µm XB-C18 100Å 150x4.60 mm (Phenomoenex 00F-4605-E0) with precolumn (AJ0-9000); solvent A: 50 mM $NH_4OAc$ pH 6.7; solvent B: 50 mM $NH_4OAc$ pH 6.7 in 70% MeCN; elution program: from 0 to 35% B in25 min, from 35 to 60% B in 2 min, from 60 to 0% B in 3 min, hold 0% for 2 min; flowrate: 1.1 ml/min. Product ret. time ~ 20.6 min.

Conversion of amino-11-UTP to Cy3B-11-UTP was similar (90–92%) for both of reaction conditions (pH 7 and 9). Collected fractions were combined freeze-dried two times (second after resuspension in MQ water) to remove the excess of ammonium acetate.

Combined products after freeze-drying were dissolved in 2.00 ml of MQ water. The concentration of the afforded solution was measured with nanodrop (C = 74.4 µM, $A_{289}$ = 0.127, $A_{559}$ = 0.901, 1 mm optical pathway, $E_{559}$= 121 $mM^{-1}cm^{-1}$). The solution was freeze-dried one more to afford the Cy3B-UTP product as a magenta solid (149 nmol, yield 78%)

The solid was dissolved in ~50 µl of MQ water for further experiments (C = 2.93 mM, $A_{560}$ = 35.4, 1 mm optical pathway, $E_{559}$= 121 $mM^{-1}cm^{-1}$). Sample of the products was analyzed with MS(-): m/z = 1191.2996, 595.1334, 396.4234; expected m/z(-): $(M-H)^{-}$ = 1191.2594, $(M-2H)^{2-}$ = 595.1260, $(M-3H)^{3-}$ = 396.4149). Mass spectrometry analysis was performed using an Agilent 1100 coupled to a microTOF mass spectrometer (Bruker Daltonik). The microTOF mass spectrometer (Bruker Daltonik) was operated in positive ion mode over an m/z range of 100–1300, and MS1 scans were recorded. Data processing was conducted using Compass™ 1.7 DataAnalysis 4.2 software (Bruker Daltonik). Spectra were deconvoluted using the MaximumEntropy algorithm with an instrument resolving power of 10,000.

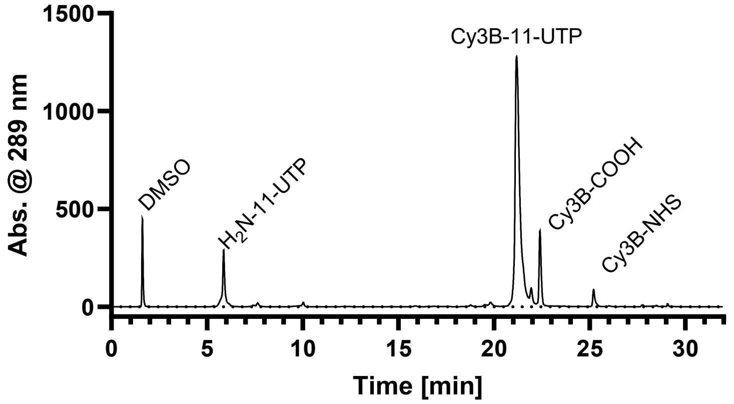

HPLC profile of the reaction mixture (cond. pH 9)

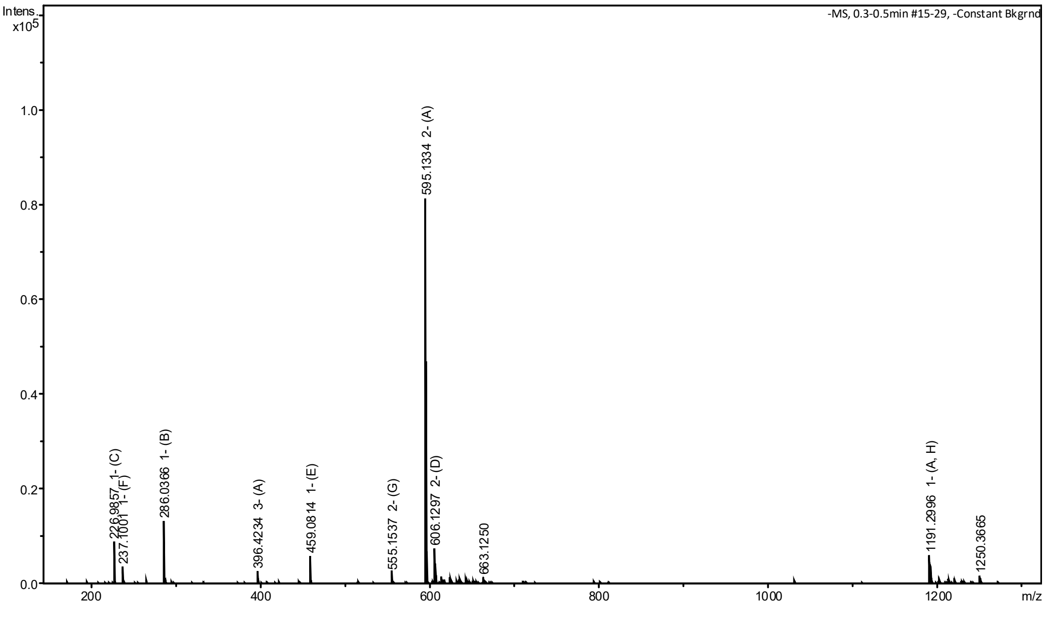

MS spectrum of the final product

#### T7 RNAP Alexa Fluor 488 maleimide

Alexa Fluor™ 488 C5 maleimide (ThermoFischer Scentific) was used for maleimide labelling of T7 RNAP. No specific tag was used due to internal cysteine residues of T7 RNAP. The protein was adjusted to a concentration of 40 µM and incubated overnight with 25 µg dye dissolved in DMSO at 4 °C. Free dye was separated by a gravity flow column. A labelling efficiency of 168 % was determined. Fractions were pooled and stored at – 80 °C. (Storage buffer: 50 mM Tris-HCl, 100 mM NaCl, 20 mM β-ME, 1 mM EDTA, 50% Glycerol, 0.1% (w/v) Triton® X-100, pH 7.9 @ 25°C.)

#### Preparation of DNA templates

5'-Modified DNA templates (look List of DNA templates products and sequences section) were prepared with PCR with Phusion DNA polymerase (Thermo Fisher), a plasmid template (look Plasmid maps and sequences, section) and sequence-specific primers (look List of oligonucleotides section). Reactions (8 x 50 µl, 400 µl in total) contained 80 ng plasmid template, 0.25 µM primers, 1x Phusion HF buffer, 0.2 mM dNTPs, and 8 U Phusion polymerase, assembled in 0.5 ml thin-walled PCR tube strip.

Immediately after assembly at room temperature, the samples were run in the following thermocycler program: 98 °C for 5 min; 35 cycles of 98 °C for 20 s, 60 °C for 20 s, 72 °C for 40 s; and 72 °C for 10 min. The individual reactions were pooled and DpnI FD (Thermo Fisher) was added (1 µl per 400 µl PCR) and incubated at 37 °C for 30 min to digest template DNA.

PCR products were purified with the PureLink Quick PCR Purification Kit (Thermo Fisher) according to the manufacturer's protocol. DNA was eluted in 2 × 50 µl 10 mM Tris-HCl, pH 8.5, and quantified with NanoDrop. Product size and integrity were verified by agarose gel electrophoresis (0.8% agarose in TAE), loading ~50 ng DNA and a GeneRuler 1 kb Plus DNA Ladder (Thermo Fisher) at 100 V for 45 min, followed by fluorescence imaging (GE Healthcare Amersham Imager 600). The imaging was repeated after gel staining in SybrSafe (Invitrogen).

#### RNA transcription in bulk

In vitro transcription was performed using T7 RNA Polymerase HC (Thermo Fisher) and PCR-derived DNA templates. Reactions were assembled in 1.5 ml RNase-free tubes and mixed thoroughly at room temperature to avoid precipitation. A typical reaction mixture (60 µl) contained transcription buffer (20 mM Hepes-KOH pH 8.0, 10 mM NaOAc, 5 mM $Mg(OAc)_2$, 2 mM spermidine, 2 mM DTT), ATP (0.25 mM), GTP (0.25 mM), UTP (0.25 mM), CTP (0.25 mM), Cy3B-UTP (2.5 µM), DNA template (900 ng), Ribolock RNase inhibitor (0.25 U/µl, Thermo), Inorganic pyrophosphatase (0.5 U/µl, Thermo). This mixture (30 µl, containing 0.9 µg of DNA).

Reactions were incubated at 37 °C for 2 h. Residual DNA was removed by adding 0.5 µl of DNase I (Thermo Fisher) incubating 30 min at 37 °C. RNA was purified using the Monarch RNA Clean-up Kit 10 µg (NEB) according to the manufacturer's instructions, with elution in RNase-free water.

RNA quality and approximate size were assessed by agarose gel electrophoresis (1% agarose in TBE), loading ~100 ng RNA together with RiboRuler HR (Thermo Fisher) at 80 V for 45 min, followed by fluorescence imaging (GE Healthcare Amersham Imager 600). The imaging was repeated after gel staining in SybrSafe (Invitrogen).

#### Preparation of supported lipid bilayers (SLBs)

Small unilamellar vesicles (SUVs) were prepared from lipids mixtures dissolved in chloroform. First chloroform lipid stocks were combined in glass vials, and the mixtures were dried under a stream of nitrogen to form thin lipid films. The films were briefly rehydrated in chloroform and dried again to improve homogeneity, after which they were subjected to an vacuum-drying for one hour to remove residual solvent. The dried lipid films were then rehydrated in 40 mM tris-HCl pH 7.5 (4 mg/ml total lipid concentration) at room temperature, and the suspensions were vortexed and pipetted repeatedly until the multilamellar lipid material fully detached from the glass surfaces and formed a uniform dispersion. For lipids with higher phase-transition temperatures (i.e. DPPC), the rehydration step was carried out at elevated temperature (i.e. 50°C) to ensure complete swelling of the films. The resulting multilamellar vesicle suspensions were divided into 20 µl aliquots in 200 µl PCR tubes and frozen (-20°C) for storage.

Glass coverslips (24x50 mm, 1.5 thickness) for membrane deposition were prepared in parallel. Coverslips were cleaned by sequential rinsing with water, ethanol, and isopropanol, followed by $O_2$ plasma treatment (Diener Electronic Zepto instrument, for 30 seconds at 0.3 mbar and 50% power) to remove remaining contaminants and render the surface hydrophilic. Removable multiwell chambers (Culture-Insert 3 Well in µ-Dish 35 mm high, Ibidi 80366) were affixed to the cleaned coverslips.

For SUV production, aliquots were thawed, briefly mixed, and subjected to bath sonication at room temperature (with heating for DPPC) until the suspensions became optically clear. SUVs were used immediately to avoid instability during storage.

Supported lipid bilayers (SLBs) were prepared by depositing small unilamellar vesicles onto freshly prepared chambers. All steps were carried out on a thermoblock surface and thermally equilibrated buffers at 40 °C (50 °C for DPPC).Vesicles (20 µl) were mixed with deposition buffer (140 µl, 20 mM Hepes-KOH pH 8.0, 10 mM NaOAc, 5 mM $Mg(OAc)_2$) and transferred to the chambers, followed by a incubation for 5 min to promote fusion and bilayer formation. During incubation (after ~3 min), a 0.6 µl of 100 mM $CaCl_2$ solution was added to facilitate vesicle rupture. After incubation, the bilayers were gently and sequentially washed with 100 µl of IVT buffer (20 mM Hepes-KOH pH

8.0, 10 mM NaOAc, 5 mM $Mg(OAc)_2$, 2 mM spermidine, 2 mM DTT), and twice with 100 µl of wash buffer (20 mM Hepes-KOH pH 8.0, 10 mM NaOAc) to remove unfused vesicles and residual salts. The chambers were covered with a glass coverslip (18x18 mm) to avoid evaporation and kept at 40°C until the start of the experiment (within next two hours).

#### Preparation of membrane-bound transcription (MBT) set-up

Following SLB formation, the buffer environment was gradually exchanged to the transcription buffer (performed at the thermoblock as described in the SLB preparation section). Chambers were filled with wash buffer (110 µl of total liquid volume in a chamber) and partially aspirated (removal and subsequent addition of 50 µl of liquid) in a stepwise manner, first with wash buffer and subsequently with buffers of increasing ionic and cofactor strength (IVT x1, and IVT x2 buffers), ensuring passive mixing and leaving a defined final volume (30 µl). This established the reaction environment for in vitro transcription on the membrane surface (final buffer composition: 20 mM Hepes-KOH pH 8.0, 10 mM NaOAc, 5 mM $Mg(OAc)_2$, 2 mM spermidine, 2 mM DTT). Such chamber was transferred to a thermally equilibrated microscope environmental chamber (37°C) and covered with a glass coverslip (18x18 mm) to avoid evaporation.

The transcription mixture lacking RNA polymerase was assembled separately and pre-incubated at 37°C. A typical reaction mixture contained transcription buffer (20 mM Hepes-KOH pH 8.0, 10 mM NaOAc, 5 mM $Mg(OAc)_2$, 2 mM spermidine, 2 mM DTT), ATP (0.5 mM), GTP (0.5 mM), UTP (0.5 mM), CTP (0.5 mM), Cy3B-UTP (5 µM), DNA template (30 ng/µl), Ribolock RNase inhibitor (0.5 U/µl, Thermo), Inorganic pyrophosphatase (1 U/µl, Thermo). This mixture (30 µl, containing 0.9 µg of DNA) was then introduced dropwise into the center of each chamber to avoid perturbing the bilayer and covered with a glass coverslip to avoid evaporation. Samples were allowed to equilibrate within the microscope environmental chamber for one hour prior to addition of T7 RNA polymerase.

T7 RNA polymerase (200 U/µl, Thermo) was diluted with transcription buffer to obtain a series of working concentrations (typically 24 U/µl). The enzyme solution (5 µl) was added to the center of each chamber and gently mixed by pipetting to start the reaction.

After the reaction, reaction mixtures were collected from each chamber for downstream RNA purification and analysis (as described in Transcription in bulk section).

For different experimental applications, SLBs and MBT samples were prepared with other types and sized of the chambers. All prepared according to abovementioned protocol, but with adjustments to all liquid volumes and total amount of DNA by scaling to the total sample volume and the SLB area, respectively:

| Chamber type | Catalog No. | Total sample volume [µl] | SLB area [$cm^2$] | DNA used [µg] |
|---|---|---|---|---|
| Culture-Inserts 3 Well | Ibidi 80369 | 60 | 0.22 | 0.18-1.8 |
| 12 well chamber | Ibidi 81201 | 300 | 0.56 | 1.5 |
| 3 Well Chamber | Ibidi 80381 | 1000 | 1.66 | 5 |

#### Total internal reflection fluorescence (TIRF) microscopy

Total internal reflection fluorescence (TIRF) imaging was performed on a Zeiss Elyra 7 microscope using an alpha Plan-Apochromat 63x/1.46 Oil Korr M27 Var2 oil-immersion objective (Carl Zeiss). Atto 488 was excited using a 488 nm diode laser; Cy3B using a 561 nm diode laser; and Alexa 647N using a 642 nm diode laser. The optical path consisted of a LBF 405/488/561/642 main beam splitter and a LP 560 beam splitter. Images were taken at 512 × 512 px at a pixel scaling of 0.098 µm × 0.098 µm, using an exposure time of 50 ms.

Time-resolved TIRF imaging was performed prior to and immediately after transcription initiation. Single-frame images and tiled acquisitions were captured at defined intervals throughout the reaction. To remove imaging artefacts due to particle interference in the detection path, TIRF images were processed with a pseudo flat-field background correction using the BioVoxxel plugin in ImageJ/Fiji. (1)

#### Confocal fluorescence microscopy

Confocal imaging was performed at a Zeiss LSM980 confocal laser scanning microscope using a Zeiss C-Apochromat 40×/1.20 water-immersion objective (Carl Zeiss). Atto 488 was excited using a 488 nm diode laser; Cy3B using a 561 nm diode laser; and Alexa 647N using a 642 nm diode laser. The optical path for 488 channel consisted of a 488/561 main beam splitter; and for 561 and 640 channels, a 488/561/639 main beam splitter. Channels were recoded separately to avoid cross-talk. The temperature of the built-in incubation unit was kept at 37 °C during imaging.

#### Fluorescence recovery after photobleaching (FRAP)

Confocal laser scanning FRAP was performed at the Zeiss LSM980 microscope described above. Circular region of interests (ROIs) measuring 5.0 µm in diameter were defined for photobleaching. FRAP recordings implemented a pinhole size of 1.0 Airy unit, a pixel number of 512 × 512 at a pixel scaling of 0.083 µm × 0.083 µm, and a pixel dwell time of 0.34 µs.

#### Spatial cross-correlation analysis

Spatial cross-correlation analysis was performed using the JaCoP (Just Another Colocalization Plugin) plugin in ImageJ/Fiji. (2) TIRF images were acquired and background-corrected as described above. Spatial cross-correlation between imaging channels was analysed with a maximum shift of 20 px. The cross-correlation results were used to assess phase separation through co- or anti-localization of the imaging channels during the course of transcription.

#### Number and brightness (N&B) analysis

Number and brightness analysis was conducted by measuring the pixel-wise fluorescence intensity and its fluctuations to estimate the apparent molecular number and brightness of fluorescent species within the imaged evanescent field. Pixel dwell times are set to be small compared to the molecular transit times to capture the effect of oligomerization. The resulting brightness maps were used to assess RNA aggregation during transcription.

#### Atomic force microscopy (AFM)

Atomic force microscopy was performed on a JPK Nanowizard 3 in QI (quantitative imaging) mode using BL-AC40TSC2 (Olympus) cantilevers. The samples were imaged in transcription buffer x1 (). The set point force was 0.15–0.2 nN, acquisition speed 62.5 µm $s^{-1}$, Z range 5 µm and the Z length was 150 nm. Images were processed in Gwyddion (v2.58, http://gwyddion.net/). The image processing included plane leveling, polynomial row alignment, scar correction, and conservative denoising.
